# Neural representation of action symbols in primate frontal cortex

**DOI:** 10.1101/2025.03.03.641276

**Authors:** Lucas Y. Tian, Kedar U. Garzón Gupta, Daniel J. Hanuska, Adam G. Rouse, Mark A. G. Eldridge, Marc H. Schieber, Xiao-Jing Wang, Joshua B. Tenenbaum, Winrich A. Freiwald

## Abstract

A hallmark of intelligence is proficiency in solving new problems, including those that differ dramatically from problems seen before. Problem-solving, in turn, depends on goal-directed generation of novel ideas and behaviors^1^, which has been proposed to rely on internal representations of discrete units, or symbols, and processes that recombine these units into a large set of possible composite representations^1–9^. Although this view has been influential in formulating cognitive-level explanations of behavior, definitive evidence for a neuronal substrate of symbols has remained elusive. Here, we identify a neural population encoding action symbols—recombinable representations of discrete units of motor behavior—localized to a specific area of frontal cortex. In macaque monkeys performing a drawing-like task, we found behavioral evidence that action elements (strokes) exhibit three critical features indicating an underlying symbolic representation: (i) invariance over low-level motor parameters; (ii) categorical structure, reflecting discrete types of action; and (iii) recombination into novel sequences. In simultaneous neural recordings across motor, premotor, and prefrontal cortex, we found that planning-related population activity in ventral premotor cortex (PMv) encodes actions in a manner that, like behavior, reflects motor invariance, categorical structure, and recombination. Activity in no other recorded area exhibited these three properties of symbols. These findings reveal a neural representation of action symbols localized to PMv, and therefore identify a putative neural substrate for symbolic cognitive operations.

## Introduction

Understanding the mechanisms of intelligence requires explaining generalization, especially to situations or problems that differ considerably from those previously encountered. For example, if asked to “draw an animal that doesn’t exist”, children can generalize from prior experience to produce an imaginary animal, such as a dog-like creature with six legs, three camel humps, and three pig tails^10^. An influential hypothesis is that such generalization depends on an internal representation of discrete units (symbols) that can be recombined into composite representations, in a process called compositional generalization^1–9^. Symbols enable systematic derivation of novel representations from a few reused components, such as new animals constructed from known parts (*animal = 1 torso + 8 arms + 4 legs*). This hypothesis is not restricted to concepts explicitly represented as symbol systems in language, computer programs, and mathematics, but applies broadly even to abilities not superficially symbolic^5,7,9^, including, in humans, geometry^7^, handwriting^11^, drawing^12^, dance^13^, musicianship^7^, and speech^14^, and, also in non-human animals, reasoning (logical^9^, spatial^15,16^, physical^9^, numerical^17^, and social^18^), artificial grammar learning^19–21^, and communication^22,23^.

Despite behavior suggesting symbolic representations, we lack definitive evidence for whether and how symbols are implemented in neuronal substrates. This is especially problematic given uncertainty over how symbols reconcile with prominent models of cognition that do not presuppose symbols, including those based on distributed processing in neural networks^24–26^, dynamical systems^27,28^, and cognitive maps^29–31^. Given that symbols are discrete representational units that are systematically recombined, a neural population representing symbols should exhibit three essential properties: (i) invariance, (ii) categorical structure, and (iii) recombination. Invariance means that activity is independent of variables irrelevant to the task goal. Categorical structure means the population should express one distinct activity pattern per symbol and be biased towards these discrete patterns even with continuous variation in task parameters. Recombination implies that a symbol’s activity pattern should occur in all contexts in which it is composed with other symbols.

Recordings during cognitive tasks have revealed a diversity of invariant representations, including of rules^32,33^, actions^34,35^, sequences^36,37^, numerical concepts^8^, perceptual categories^38^, visual objects^39^, cognitive maps^29,31^, and other abstract concepts^40^. These findings reveal a striking capacity for invariance, and implicate specific regions, including prefrontal cortex^32,33,36–38^ and medial temporal lobe^29,31,32,40^. However, it is unclear whether these representations exhibit the other properties expected for symbols. First, with a few exceptions^38^, prior studies did not systematically assess categorical structure by testing whether activity varies discretely with continuous variation in task parameters. Second, evidence for recombination in neural activity is also rare, with a notable exception in hippocampus: activity encoding spatial paths appears to reuse parts of previous paths in novel sequences^41,42^. However, whether these continuous paths reflect recombination of discrete components is unclear. Third, the tasks in these hippocampal studies and other studies on invariant representations did not, to our knowledge, test compositional generalization, making it challenging to determine whether and how identified activity patterns support compositional processes. Thus, we still lack evidence for a neural representation of symbols—one that jointly exhibits invariance, categorical structure, and recombination in the behavioral setting of compositional generalization.

To identify neural populations that may express these properties of symbols, we established a task involving symbol-based compositional generalization, implemented in macaque monkeys (**Fig. 1a, b**). This task involves generating novel, goal-directed actions, an ability hypothesized to involve symbolic representations in the form of discrete units of motor behavior (action symbols) that are recombined into sequences^11–14,43–46^. For example, imitating a dance by observation may depend on symbolic representations of dance poses^13^. Action symbols are also essential to various models of action sequencing, including in handwriting^11,43^, drawing^12^, object manipulation^47^, and tool use^48^, and may be related to movement segments (or “syllables”) that have been identified in naturalistic behaviors^49,50^. These studies suggest that a task involving compositional generalization in action sequencing would enable the systematic investigation of action symbols. Here, we establish such a task and then, through behavioral and neural analyses, identify a neural representation of action symbols in PMv.

**Fig. 1.**
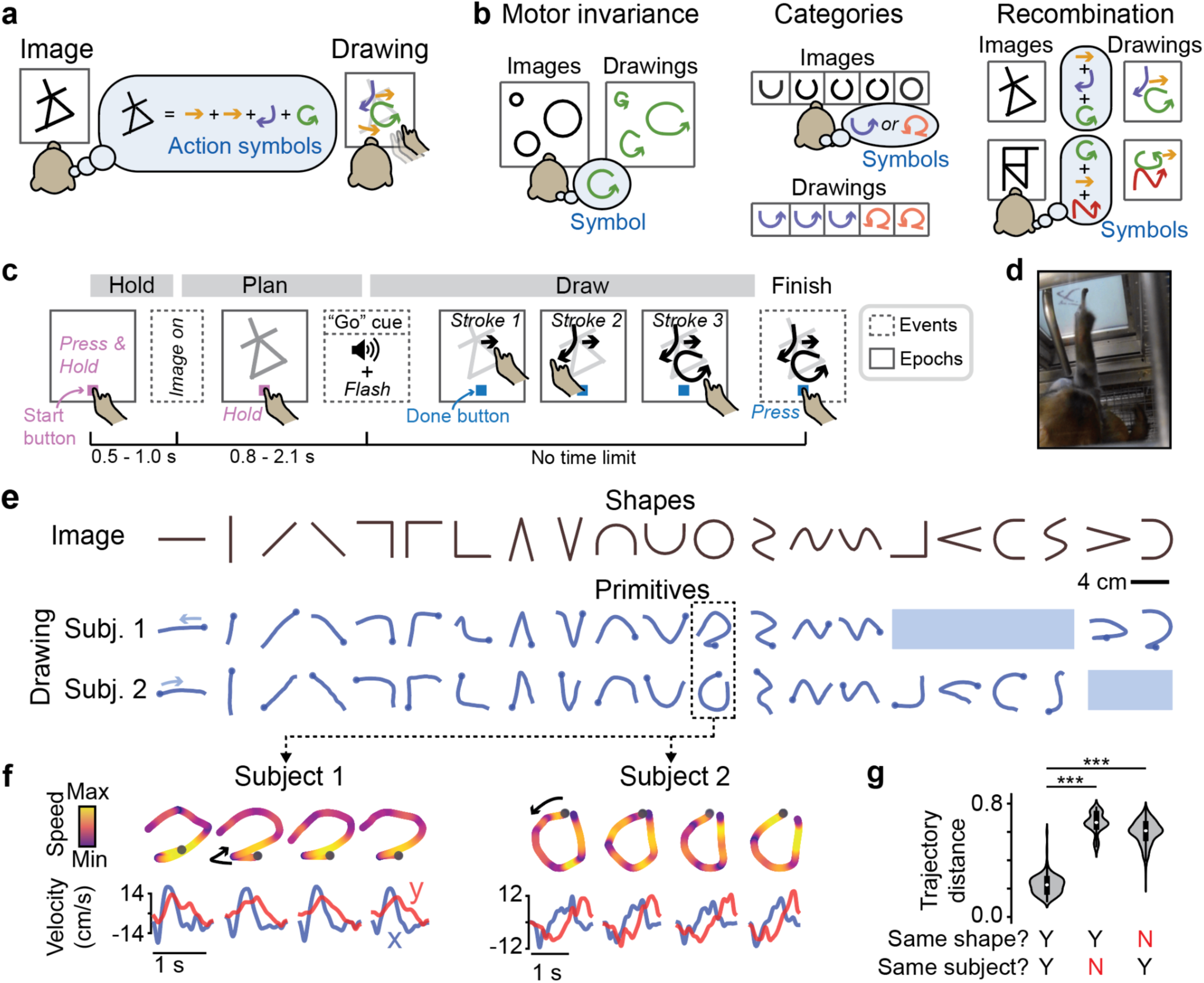
Learned stroke primitives in a drawing-like task paradigm. (a) Experimental paradigm. Subjects draw images by tracing on a touchscreen, with no cues for stroke type and order. We tested if strokes are internally represented as learned action symbols. (b) Three essential features of a symbolic representation, which we tested in behavior and neural activity: invariance, categorical structure, and recombination. Schematics depict predictions by the action symbol hypothesis. Alternative predictions in **Figs. 2a, f, k**. (c) Trial structure, showing three discrete events (dashed boxes) and sustained epochs (solid boxes). “Hold”: press and hold a “start” button, enforcing a consistent posture. “Plan”: subjects see the image but must maintain hold on “start” button. “Draw”: subjects produce strokes. Subjects can report completion any time by pressing “done”. They receive juice reward based on drawing performance (Methods). “Buttons” always refers to virtual touchscreen buttons. (d) Photograph of during in-cage training. Metal tube is reward spout. Videos in **Supplementary Videos 1-10**. (e) Learned stroke primitives (blue) for each shape (black). Stroke onsets are marked with a ball. Blue shading marks images for which subjects did not readily learn primitives, either because they used two strokes or one stroke in a variable manner (Methods). Drawings average over 15 trials. (f) Four example trials per subject for the “circle” primitive, depicted spatially (top) and temporally (bottom). (g) Summary of stroke trajectory distances for trial pairs defined along two dimensions: same shape (yes/no) and same subject (yes/no). Violin plots represent kernel density estimates, with medians and quartiles. n = 300 (“same shape, same subject”: YY and YN; 15 shapes x 10 trials x 2 subjects), n = 4200 (NY; 15 shapes x 10 trials x 14 other shapes x 2 subjects). ***, p = 6.10 x 10^-5^, two-sided Wilcoxon signed-rank test (W = 0, n = 15 shapes, averaged over trials and subjects).

## Results

### Learned stroke primitives in a drawing-like task

We developed a drawing-like task for macaque monkeys modeled after studies of human drawing^12^ and handwriting^11,43^. We trained two subjects to draw geometric figures by tracing on a touchscreen (**Fig. 1a,c,d**; **Supplementary Videos 1-10**; setup in **Extended Data Fig. 1**). Subjects drew images that varied across trials, and were rewarded for accuracy, quantified as the spatial similarity between the drawn and “target” images (primarily using the Hausdorff distance; see Methods).

Once subjects acquired the basic skills to trace images, they practiced drawing a diverse set of simple shapes, each using one stroke (**Fig. 1e**). Although we did not force subjects to use particular spatio-temporal trajectories (i.e., strokes), each subject converged on a set of strokes, one for each shape. We call this set the subject’s “primitives” (**Fig. 1e**). We found that primitives were idiosyncratic to each subject and shape. To analyze primitives, we devised a “trajectory distance” metric based on the Euclidean distance between two strokes’ normalized velocity time series (Methods). Within each subject, each shape was consistently drawn with the same primitive (examples in **Fig. 1f**; quantification in **Fig. 1g**), and different shapes were drawn with distinct primitives (**Figs. 1e-g**). A further decoding-based assessment of across-trial consistency and across-shape distinctiveness revealed that primitives were easily distinguishable from each other based on single-trial stroke trajectories (**Extended Data Fig. 2**). Primitives were also unique to each subject; the two subjects used different primitives even for identical shapes (compare primitives in **Figs. 1e, f**; quantification in **Fig. 1g**).

The finding that stroke primitives are idiosyncratic to shape and subject raised the possibility that they reflect learned action symbols. To test this, we determined how subjects generalize to draw new figures that assess three behavioral properties that, in combination, indicate a symbolic representation: motor invariance, categorical structure, and recombination (**Fig. 1b**).

### Primitives exhibit motor invariance over location and size

If subjects represent stroke primitives as symbols, then stroke primitives should exhibit motor invariance. That is, each primitive’s idiosyncratic trajectory should generalize across low-level motor parameters (e.g., muscle activity patterns), as seen in handwriting and other skills^51^. To test motor invariance, we presented each shape at varying sizes and locations, to an extent expected to elicit variation in motor cortical and muscle activity (Methods). Motor invariance predicts similar stroke trajectories across location and size (“Symbols”, **Fig. 2a**). Alternatively, if individual responses were memorized for each stimulus, then the subject would have difficulty generalizing, as seen in some inflexible skills^52^ (“Fail” in **Fig. 2a**). A third alternative strategy could prioritize efficiency by minimizing movement from where the hand starts before stroke onset, thus predicting different stroke trajectories for different locations and sizes (“Efficient” in **Fig. 2a**). We found that stroke trajectories were similar across locations and sizes (**Figs. 2b-e**), indicating that stroke primitives exhibit motor invariance.

**Fig. 2.**
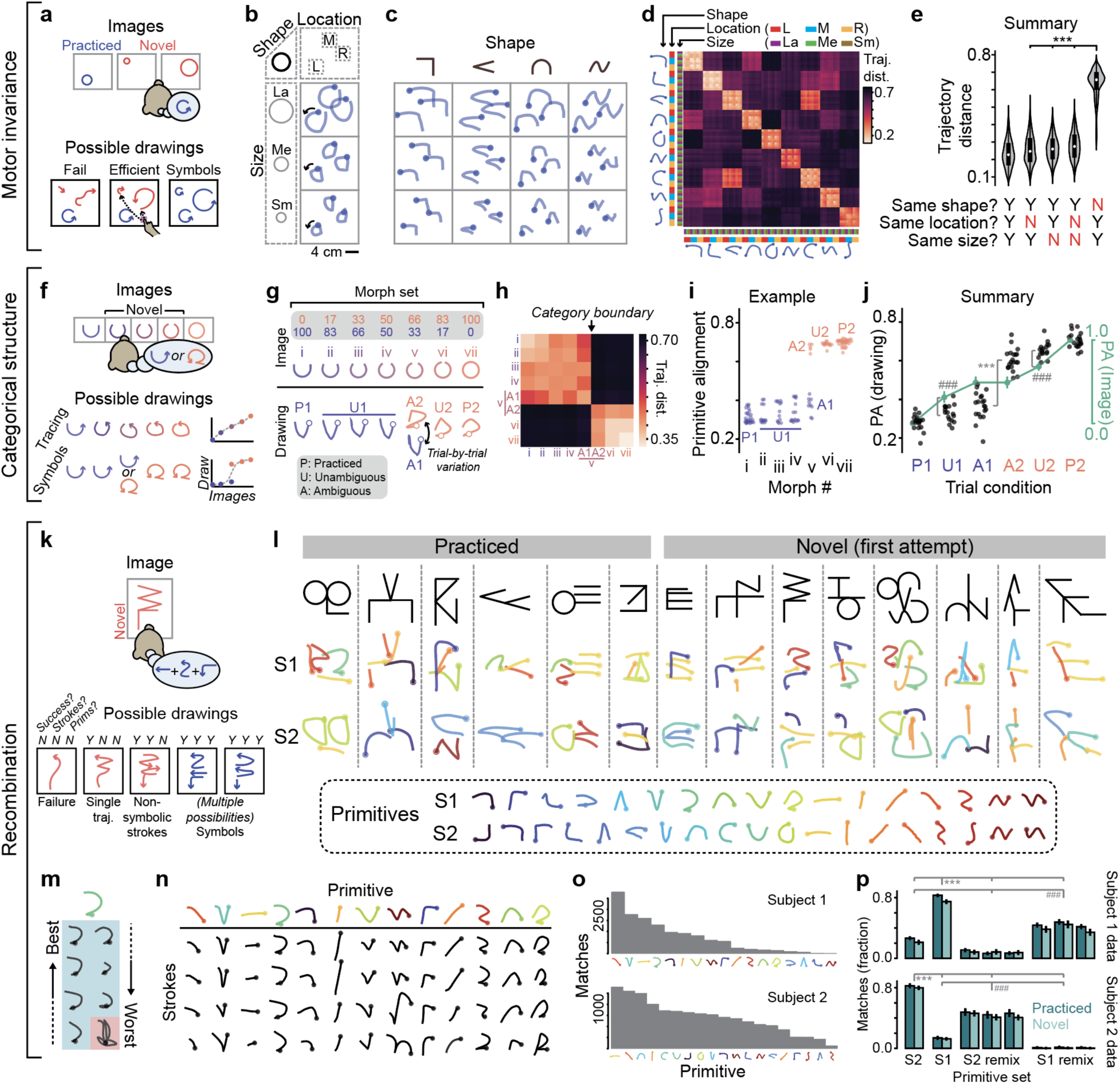
Behavioral evidence for action symbols: motor invariance, categorical structure, and recombination into sequences. (a) Experiment testing motor invariance. Shapes vary in size (novel) and location across trials (gray rectangles). Shown are three possible drawing outcomes; “symbols” reuses primitives. (b) Example drawings for “circle” shape, varying in location (left, middle, right) and size (small, medium, large). Drawings from three trials overlaid on each panel (subject 2). (c) Example drawings for more shapes. (d) Mean pairwise trajectory distance between each shape-location-size condition (n = 9 trials per condition). (e) Summary of pairwise trajectory distances, with trial-pairs defined along three dimensions: same shape (yes/no), same size (yes/no), and same location (yes/no), plotting kernel density estimates, with medians and quartiles. Each datapoint is one trial’s average distance to one shape-size-location condition. n = 648 (same shape/size/location: YYY), 1296 (YYN), 1296 (YNY), 2592 (YNN), 5184 (NYY), from a pool of 729 trials (81 conditions x 9 trials each). ***, two-sided Wilcoxon signed-rank tests comparing NYY to others: vs. YNY (W = 0, p = 5.36 x 10^-15^), vs. YYN (W = 0, p = 5.36 x 10^-15^), vs. YNN (W = 0, p = 5.36 x 10^-15^); n = 81 shape-size-location conditions). (f) Experiment testing categorical structure. Given novel images that morph between two well-practiced shapes, we consider two hypotheses, “Tracing” and “Symbols” (see main text). (g) Example morph set, varying between two well-practiced images: morph *i* (mixture ratio of 100% “U”, 0% circle) and morph *vii* (0% “U”, 100% circle). Example drawings are shown below each image. Morph *v* elicited trial-by-trial variation (A1, A2). Trials are grouped into three conditions: well-practiced (P), unambiguously one primitive (U), or ambiguous (A). (h) Pairwise trajectory distances for the example morph set in panel g. Category boundary is defined as the morph found to elicit variation in drawing. n = 5 - 30 trials (range). (i) Single-trial primitive alignment (PA) vs. morph for the morph set in panel g. n = 5 - 30 trials (range). (j) Summary of PA across experiments, using both image data (green, mean and 95% CI) and drawing trajectories (black). Each drawing datapoint represents a single morph set [n = 20 sets, (13 for subject 1; 7 for subject 2)]. Image-based scores were rescaled so endpoint values (P1 and P2) match drawing-based scores. Test for sigmoidal nonlinearity: ###, p = 1.91 x 10^-6^, two-sided Wilcoxon signed-rank test (W = 0, n = 20) that drawing < image (U1) and drawing > image (U2). Test for trial-by-trial switching: ***, p = 1.91 x 10^-6^, two-sided Wilcoxon signed-rank test (W = 0, n = 20) that A2 > A1 (drawing). (k) Experiment testing recombination in the Character task. Four possible kinds of drawings differ along three dimensions: copying success, multiple strokes, and primitive reuse (see main text). Images can be ambiguous in having multiple ways to be drawn using primitives (“symbols”). (l) Example drawings by subjects 1 and 2 given the same images. Strokes are color-coded by best match from each subject’s own primitives, which are plotted below (“Primitives”). (m) Example character strokes assigned to the primitive “reversed C” (blue background), uniformly ranging from best to worst match. The stroke with red background failed to match any primitive. Subject 1. (n) Example character strokes organized by assigned primitive, ordered from high (top row) to median (bottom row) quality match, and most to least frequent (columns, restricted to the top thirteen from distribution in panel o). Subject 1. (o) Frequency of strokes matched to each primitive. Note that the non-uniformity of this distribution is reminiscent of frequency distributions seen in language and other behaviors (although those cases may in part reflect non-uniformity in “training” experience)^53^. Note that even low-frequency primitives still passed criteria to be deemed high-quality matches (Methods). (p) Summary of evidence for primitive recombination, showing fraction of strokes (mean and 95% CI) that match a primitive, for characters performed by both subjects. Strokes (“data”) were tested against each subject’s primitive set (S1, S2) and simulated primitive sets that mix subparts of primitives into new “remixed” primitives (**Extended Data Fig. 5**). Three independent simulations are plotted separately. N (characters) = 133 (subject 1, practiced), 90 (S1, novel), 70 (S2, P), 135 (S2, N). Wilcoxon signed-rank tests: ***, comparing match to one’s own primitives vs. other. For S1 data: S1 vs. S2 (W = 0, p = 6.75 x 10^-25^), vs S1 remix (W = 125, p = 3.43 x 10^-22^), vs S2 remix (W = 0, p = 8.75 x 10^-25^); for S2 data: S2 vs S1 (W = 49, p = 6.26 x 10^-23^), vs S2 remix (W = 424.5, p = 3.61 x 10^-16^), vs S1 remix (W = 0, p = 5.82 x 10^-25^). ###, comparing one’s own remixed primitives vs. others. For S1 data: S1 remix vs S2 (W = 995, p = 1.41 x 10^-11^), vs S2 remix (W = 351, p = 1.90 x 10^-20^). For S2 data: S2 remix vs S1 (W = 731.5, p = 1.09 x 10^-11^), vs S1 remix (W = 1, p = 2.85 x 10^-20^). For remixed primitives, the least significant of the three simulations is shown. Sample sizes are identical for all tests (n = 141 characters, combining novel and practiced).

### Primitives exhibit categorical structure

If stroke primitives are represented as categorically-structured action symbols, we would expect subjects to be biased towards using their idiosyncratic primitives when challenged with new figures that interpolate, or “morph”, between learned shapes (**Fig. 2f**, “Symbols”). This would indicate that stroke primitives correspond to discrete types, similar to stroke categories in Chinese characters^43^ or phonemes in speech^14^. In contrast, if subjects simply trace images without interpreting them as action symbols, then we would expect drawings to closely match the interpolated figures (**Fig. 2f**, “Tracing”). We presented images randomly sampled on each trial by linearly morphing between two practiced shapes. Each “morph set” consisted of two practiced shapes and four to five morphs between them. We tested whether drawings reflect a categorical boundary in the subject’s interpretation of these images, which would manifest in two behavioral hallmarks (**Fig. 2f**, “Symbols”): a steep sigmoidal relationship between image and drawing variation^38^—such that images on the same side of the category boundary are drawn similarly, while images across are drawn differently—and trial-by-trial variation between two distinct primitives for images close to the boundary.

We found evidence for both hallmarks of categorical structure. In the example shown in **Fig. 2g** (more in **Extended Data Fig. 3**), the images varied in the extent to which the top was closed, ranging linearly between two practiced shapes, a “U” (morph *i* in **Fig. 2g**) and a circle (morph *vii*). We defined the category boundary as the morph that elicited trial-by-trial variation between primitives 1 and 2 (morph *v*). We found that drawings for morphs to the left of that boundary (U1) were drawn similarly to primitive 1 (P1), and drawings for morphs to the right (U2) were similar to primitive 2 (P2, **Fig. 2g**). This was evident as two blocks in a matrix of the pairwise trajectory distances between all morphs (**Fig. 2h**). To quantify this effect, we calculated a “primitive alignment” score, which quantifies the relative similarity of a given trial’s drawing to primitive 1 (alignment = 0) and primitive 2 (alignment = 1), and is defined as *d_1_/(d_1_* + *d_2_)*, where *d_1_* (or *d_2_*) is the average trajectory distance to primitive 1 (or 2) trials. First, we confirmed that applying this metric to score the images, using image distance instead of trajectory distance (Methods), produced a linear relationship with morph number (**Extended Data Fig. 3c**). In contrast, when applied to score behavior, this relationship was sigmoidal (**Fig. 2i**). This analysis also captured the trial-by-trial variation in behavior for morphs at the category boundary (morph *v*, **Fig. 2i**). These effects—nonlinearity and trial-by-trial variation—were consistent across subjects and morph sets (**Fig. 2j**). Thus, behavior indicates that subjects represent primitives as categorical stroke types.

### Primitives are recombined into sequences

If subjects represent stroke primitives as symbols, they should recombine primitives to construct multi-stroke drawings. We tested this using two tasks that challenge subjects with complex figures, including novel ones. The first Multi-shape task presented figures that combined two to four disconnected shapes, which could be drawn in any order. We reasoned that one possible strategy was to use a single trajectory that efficiently traces over all shapes with appropriately timed touches and raises to produce strokes and gaps (“Single trajectory”, **Extended Data Fig. 4a**). With this strategy, behavior would be biased to minimize the movement, or gap distance, between shapes, ignoring whether this leads to reuse of primitives. A second strategy would be to draw by recombining the learned stroke primitives, at the expense of potentially longer gap distances (“Symbols”, **Extended Data Fig. 4a**). We analyzed pairs of consecutive strokes for which these two strategies predicted different stroke trajectories (**Extended Data Fig. 4b,** top schematic). In all analyzed cases, subjects recombined primitives at the expense of longer movements in gaps, consistent with a symbolic representation (**Extended Data Fig. 4b**, bottom).

We then tested subjects on complex figures called characters. Although characters were designed by connecting multiple simple shapes, subjects were not forced to draw using primitives that matched these shape components, and characters had ambiguity in being consistent with multiple possible interpretations in terms of shape components. We considered four possible drawing outcomes (**Fig. 2k**): subjects may fail to accurately draw these novel figures (“Failure”), they may succeed using a single unsegmented trajectory (“Single trajectory”), they may use multiple strokes that are not in their set of learned primitives (“Non-symbolic strokes”), or they may successfully draw by reusing their own primitives (“Symbols”; note that, due to ambiguity, there can be multiple possible behaviors consistent with primitive reuse).

We found that drawings were consistent with a symbolic representation. Subjects successfully drew novel characters, and did so using multiple strokes (**Fig. 2l**, **Supplementary Videos 1-10**), thus contradicting “Failure” and “Single trajectory” predictions. Given the same image, the two subjects produced different drawings (**Fig. 2l**), raising the possibility that subjects reused their primitives. To directly test that, we quantified how often character strokes closely matched each subject’s own primitives. We first scored each stroke’s trajectory distance to each of the subject’s own primitives. Using those scores, we classified each stroke as either a “match” to a specific primitive (i.e., the primitive with the lowest trajectory distance) or as a “non-match” (i.e., high trajectory distance to every primitive). Thresholds for deciding this match were determined empirically for each primitive based on its trial-by-trial variation on Single-shape trials (Methods). Resulting matches are shown as examples in **Fig. 2m, n** and as a frequency distribution in **Fig. 2o**.

A large majority of strokes were classified as matches (>82% for both subjects, **Fig. 2p**), even for novel characters (>74%, **Fig. 2p**). Two control analyses tested the specificity of match towards one’s own primitives. First, we asked how often strokes matched the other subject’s primitives and found that this was rare (<21% for novel characters, **Fig. 2p**). Second, we asked how often strokes matched simulated “remixed” primitives designed to resemble one’s own primitives in their subparts (i.e., their first and second halves) but not in their entire trajectories (see simulated primitives in **Extended Data Fig. 5**). We found that strokes infrequently matched these “remixed” primitives (<43% for novel characters, **Fig. 2p**). Combined, these control analyses show that character strokes specifically resemble one’s own primitives (first analysis) in their idiosyncratic trajectories (second analysis). Thus, subjects generalize to draw novel figures by recombining their primitives (“Symbols” in **Fig. 2k**).

### Multi-area recordings across frontal cortex

The behavioral findings so far suggest that stroke primitives have an underlying symbolic representation. We next searched for neural correlates of these putative representations. We recorded broadly and simultaneously neurons across multiple areas of frontal cortex, using chronically implanted multi-electrode arrays (sixteen 32-channel arrays, **Fig. 3a, b**, and **Extended Data Fig. 6**). We targeted regions associated with motor, planning, and other cognitive functions (Methods), including (but not limited to) the planning and control of movements in primary motor (M1), dorsal premotor (PMd), and ventral premotor cortex (PMv)^34^ (including in handwriting^54,55^); abstraction and reasoning in dorsolateral and ventrolateral prefrontal cortex^32,36–38^ (dlPFC and vlPFC) and frontopolar cortex^56^ (FP); and sequencing in the supplementary motor areas^57^ (SMA and preSMA). We recorded 48.4 +/- 19.9 units per area (mean and S.D.) for subject 1 and 48.0 +/- 16.0 for subject 2.

**Fig. 3.**
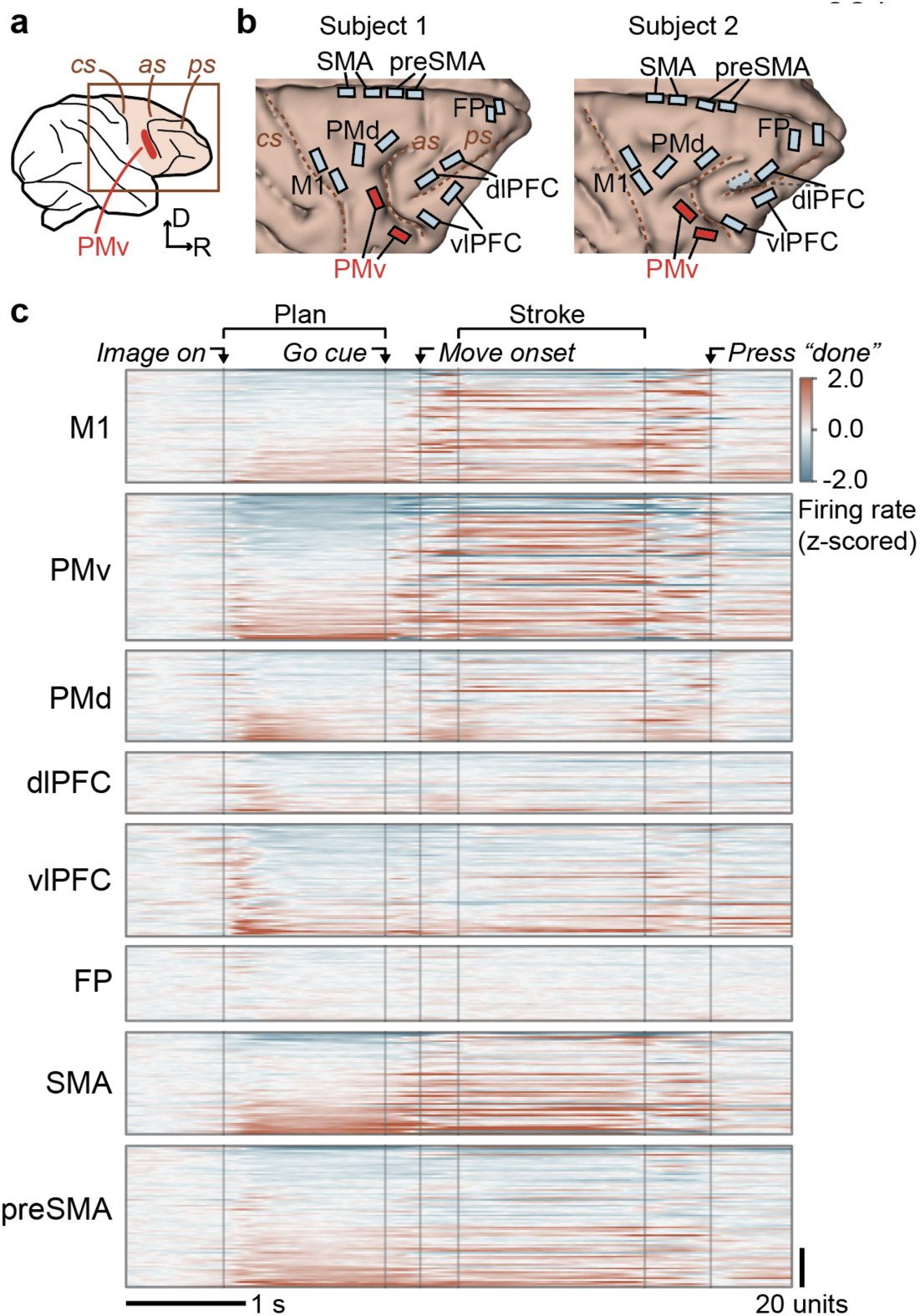
Multi-area neuronal recordings across frontal cortex. (a) Recordings were targeted to right frontal cortex, contralateral to the drawing hand. cs, central sulcus; as, arcuate sulcus; ps, principal sulcus. R, rostral. D, dorsal. (b) Arrays to scale on 3D rendering of brain. SMA and preSMA electrodes were angled to target the medial wall (**Extended Data Fig. 6**). The caudal dlPFC array for subject 2 malfunctioned. (c) Average activity across trials, grouped by brain area (panels) and split by unit (inner rows; combining single- and multi-unit). Trials were aligned by linear time-warping (Methods). Activity was z-scored relative to baseline (preceding image onset) and units were sorted by activity during planning. Subject 2.

We found clear task-related activity in all recorded areas except FP. This is evident as non-zero baseline-subtracted trial-aligned activity in **Fig. 3c**. Different areas exhibited grossly different trial-related activity patterns. For example, many units in prefrontal (vlPFC, dlPFC) and premotor (PMd, PMv) areas had rapid responses locked to image onset (**Fig. 3c**) and varied activity patterns during the planning epoch. In contrast, motor area M1 had relatively strong movement-related activity during the stroke epoch (**Fig. 3c**). Paralleling behavior, we next tested whether activity in each area encodes primitives in a manner exhibiting motor invariance, categorical structure, and recombination. We found that each of these properties was strongest in a single area, ventral premotor cortex (PMv).

### Motor-invariant encoding of primitives in PMv

In the Single-shape task, we analyzed activity during the planning epoch (between image onset and go cue) as subjects drew primitives that varied in location (**Fig. 2a-e**). PMv activity varied depending on the planned primitive, with relatively little influence of location. For example, the unit shown in **Fig. 4a, b** fired strongest for the “rotated L” primitive (black) and second strongest for the “sideways V” (blue), and this pattern repeated across locations. We visualized population activity in a two-dimensional linear projection (a primitive-encoding subspace; Methods), revealing strong variation in PMv activity due to primitive, and minimal variation due to location (**Fig. 4c,d**). For example (**Fig. 4d**), within each location (subpanel), primitive trajectories (each color) separated after image onset (“x” icon), but across locations, trajectories were similar. As a contrast, in a different area implicated in executive function, dlPFC, simultaneously-recorded activity reflected location, with minimal effect of primitive (**Fig. 4e-h**). Location-invariant encoding of primitives was apparent in PMv even in individual trials (**Extended Data Fig. 7**).

**Fig. 4.**
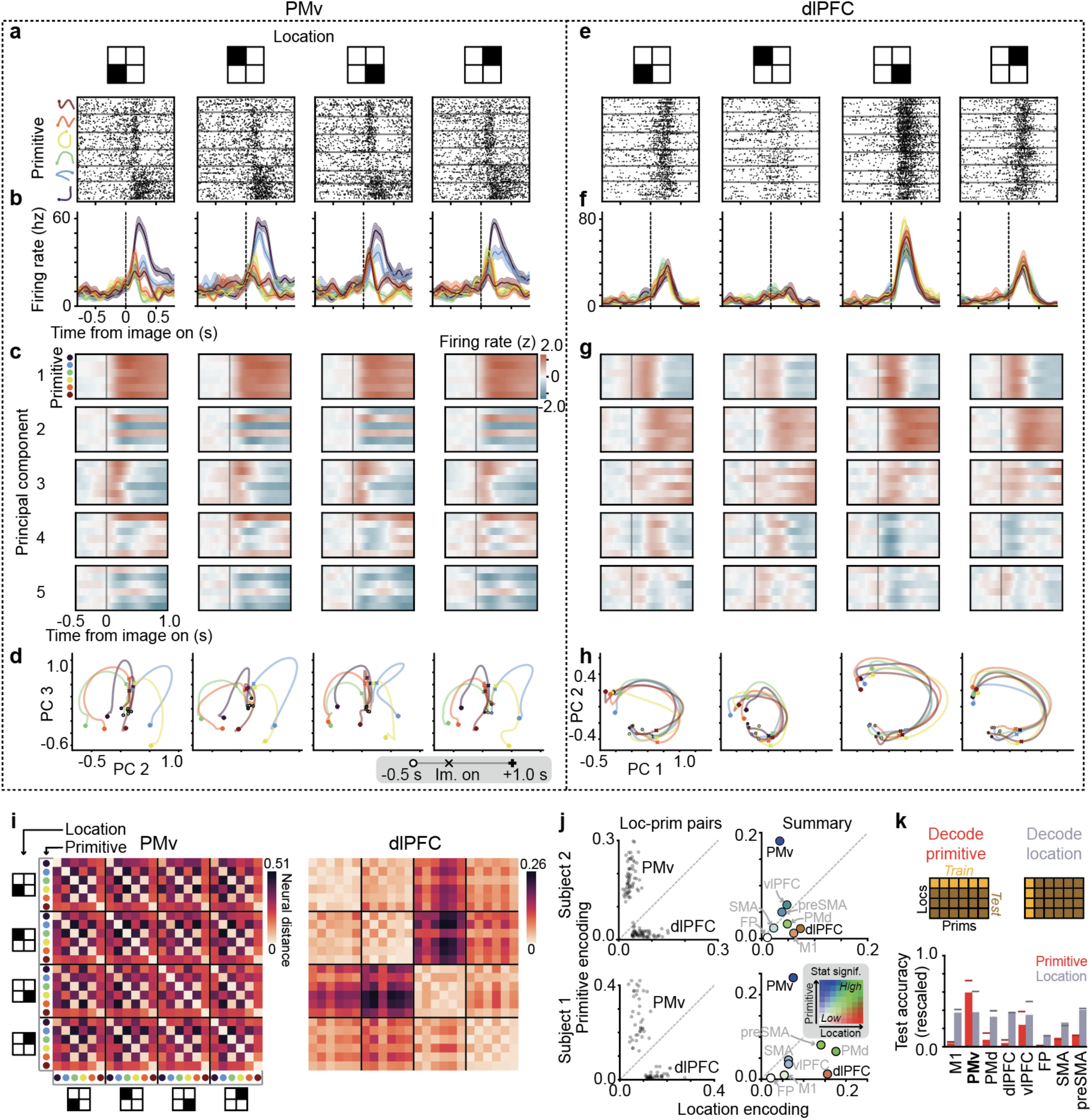
Location-invariant encoding of stroke primitives in PMv (for size-invariance see Extended Data Fig. 10). (a) Raster plot for an example PMv unit. Trials (inner rows) are grouped by primitive (outer rows) and location of drawing (columns). The finger was held on the fixation button throughout the plotted time, which is before the go cue (occurring at 1.1 - 1.5 s). (b) Trial-averaged firing rates (mean and SE) for the example PMv unit (n = 13 - 20 trials per primitive-location combination). (c) PMv population activity in primitive-encoding principal components (outer rows), showing average activity for each primitive (inner rows). Here, activity was z-scored and baseline subtracted relative to before image onset. (d) Average trajectory of PMv population activity for principal components 2 and 3. Gray legend depicts timeline of trajectories, with image onset marked with “x”. (e) Analogous to panel a, but for a simultaneously recorded dlPFC unit. (f) Analogous to panel f, but for the example dlPFC unit. (g) Analogous to panel c, but for dlPFC, plotted for PCs 1 and 2 to emphasize the main effect of location in dlPFC. (h) Analogous to panel d, but for dlPFC. (i) Pairwise neural distances between primitive-location conditions, for the session in panels a-h, averaged over time (0.05 to 0.6 s). n = 13 - 20 trials per primitive-location condition. (j) Summary of primitive and location encoding across areas and sessions. In the left two panels, points represent individual primitive-location conditions averaged over all other conditions (Methods). n = 37 (subject 1, aggregating 2 sessions), 59 (subject 2, 3 sessions). In the right two panels, each point depicts the mean. Color denotes statistical significance, indicating the number of other areas this area “beats” in pairwise statistical tests of primitive encoding and location encoding (represented in the inset heatmap, “low” to “high” ranges from no (0) to all (7) other regions “beaten”) (Methods). For example, PMv is deep blue because it has higher primitive encoding than every other area, but not higher location encoding. For statistics, each data point was a unique pair of primitive-location conditions pooled across sessions (2 for subject 1; 3 for S2). Primitive encoding (same location, different primitive): n = 93 (S1), 288 (S2). Location encoding (same primitive, different location): n = 114 (S1), 132 (S2). For exact effect sizes and p-values, see **Extended Data Fig. 8a,b**. (k) Across-condition decoder generalization for primitive (red) and location (gray). Top: schematic of train-test split method. Test accuracy is rescaled between 0 (chance) and 1 (100%). Horizontal lines represent within-condition decoding.

To quantify encoding of primitives and location, we devised a “neural distance” metric, which reflects activity dissimilarity between sets of trials representing two conditions (e.g., two location-primitive conditions). Neural distance is the average pairwise Euclidean distance between all across-condition trials, normalized by average within-condition Euclidean distance, and ranges between 0 (no difference) and 1 (a ceiling, defined as the 98th percentile of pairwise Euclidean distances). We found that neural distances in PMv were low for the same primitive across locations, visible as off-diagonal streaks in a matrix of pairwise distances (**Fig. 4i**). In contrast, the analogous matrix for dlPFC had four blocks along the diagonal, indicating similar activity within location, regardless of primitive (**Fig. 4i**). Summarizing these effects, in PMv neural distances between primitives controlling for location (“primitive encoding”) were higher than distances between locations controlling for primitives (“location encoding”) (**Fig. 4j**, left column). In dlPFC the opposite was true: primitive encoding was low and location encoding was high (**Fig. 4j**, left column). A comparison of all areas (**Fig. 4j**, right column) showed that PMv was the only one to exhibit strong primitive encoding and weak location encoding.

As further evidence that PMv’s population code for primitives is similar across locations, we measured the performance of a linear decoder, trained to decode primitives at one location, in generalizing to held-out locations^32^. Cross-location decoder generalization was strong in PMv and weaker in other areas (**Fig. 4k**).

We performed two other tests of motor invariance. First, during the initial reaching movement between the go cue and the start of drawing (**Extended Data Fig. 9a**), an epoch during which motor cortical areas are known to encode reach direction^58,59^, we found location-invariant encoding of primitives in PMv, while other motor and premotor areas (M1, PMd, and SMA) showed a mixture of primitive- and location-encoding (**Extended Data Fig. 9**). Second, PMv population activity during planning encodes primitives in a manner also invariant to size (**Extended Data Fig. 10**). Thus, PMv represents primitives in a manner invariant to the motor parameters of location and size.

### Categorical encoding of primitives in PMv

We previously found sigmoidal variation in behavior even with linear variation in images (**Figs. 2f-j**). Combining this behavioral paradigm (**Fig. 5a**) with recordings, we found that PMv activity during planning diverges towards separate primitive-representing states matching what primitive the subject will draw (**Fig. 5**). After image onset (“x” icon, **Fig. 5b**), population activity for trials in which the subject is planning to draw primitive 1 (morphs *i-iv*) separated from trials for primitive 2 (morphs *vi, vii*), such that activity diverged to two states representing primitives 1 and 2. Similarly, for the ambiguous image at the category boundary (morph *v*), trajectories diverged towards these two states depending on whether the subject was planning to draw primitive 1 or 2 (A1 or A2, in **Fig. 5c**).

**Fig. 5.**
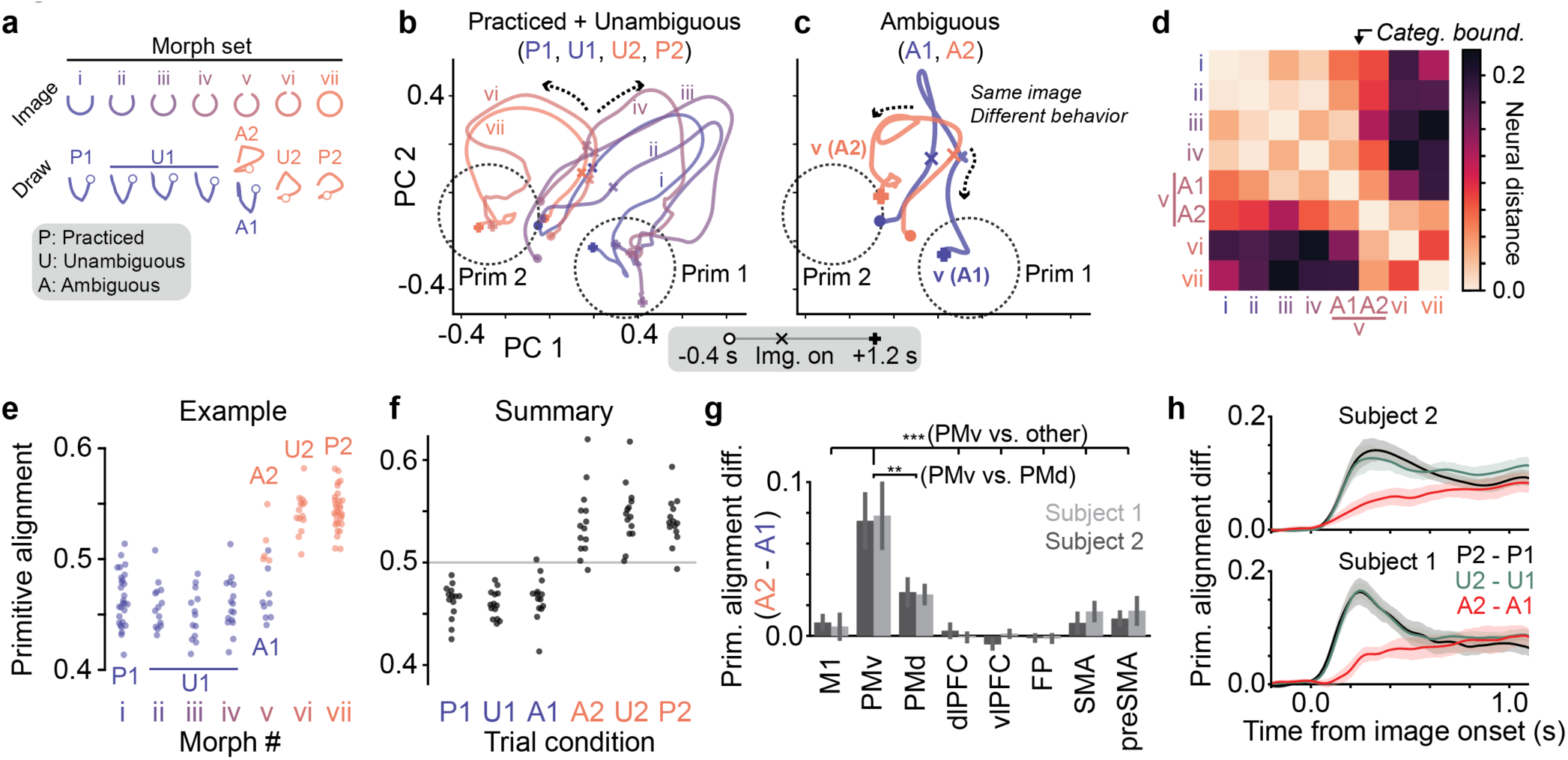
Categorical encoding of stroke primitives in PMv. (a) Schematic of example experiment. See behavior in Fig. 2f-i. (b) PMv population trajectories (description in main text). Data precede the “go cue” (occurring at 1.2 - 1.6 s). Trajectory legend is shown at bottom. Mean over n = 14 - 30 trials per trajectory. (c) PMv population trajectories for morph *v*, colored by whether the subject will draw primitive 1 (A1) or 2 (A2), overlaid on the same PCs as in panel b. n = 9 (A1) and 5 (A2) trials. (d) Pairwise neural distances, using PMv activity (n = 5 - 30 trials). (e) PMv primitive alignment for the example experiment. Datapoints reflect single trials (n = 5 - 30). (f) Summary of PMv primitive alignment vs. trial condition. Each morph set contributes one point to each condition [n = 14 morph sets (5 from S1, 9 from S2), including only the 16/20 sets that have all six trial conditions]. (g) Summary of trial-by-trial variation in primitive alignment for ambiguous images (A2 - A1), plotting mean +/- SE. **, p < 0.005; ***, p < 0.0005, paired t-tests [n = 20 morph sets, combining subjects 1 (7) and 2 (13)]. (h) Average time course of difference in primitive alignment of PMv activity, split by trial condition, and aligned to image onset (mean +/- SE). Sample size same as panel f.

We quantified this separation into two primitive-representing states by first computing the Euclidean distance between each pair of trials, and then scoring each trial with its primitive alignment [*d_1_/(d_1_* + *d_2_)*, where *d_1_* (or *d_2_*) is the average Euclidean distance to primitive 1 (or 2) trials] (**Fig. 5a**). This revealed the same two hallmarks of categorical structure we saw in behavior: sigmoidal nonlinearity and trial-by-trial switching. Sigmoidal nonlinearity is evident in the matrix of pairwise neural distances as two main blocks separated by the category boundary (**Fig. 5d**), and in the plot of primitive alignment versus morph number (**Fig. 5e, f**). Trial-by-trial switching between primitive-representing states for ambiguous images was evident in the matrix of pairwise neural distances (**Fig. 5d**, A1 is closer to morphs *i*-*iv*, while A2 is closer to morphs *vi*-*vii*), and in the primitive alignment scores (morph *v* in **Fig. 5e**; A1 vs. A2 in **Fig. 5f**). Comparing across areas, this effect was strongest in PMv (**Fig. 5g**).

We considered two neural underpinnings of the trial-by-trial switching. One possibility is that primitive choice is biased by the primitive-representing state that activity happens to be closer to at baseline, before image onset. Arguing against this, the proximity of baseline activity to primitive 1 or 2 states did not predict the chosen primitive (**Fig. 5h**, before image onset, the difference A2 - A1 is not significantly larger than zero). Another possibility is competition between states after image onset, with the winning state determining primitive choice, a possibility supported by theoretical modeling^60^ and evident in PFC^61^. Consistent with this, activity separation was slower for ambiguous images (“A2 - A1” in **Fig. 5h**), and so was behavioral reaction time (**Extended Data Fig. 11**)^60^.

Categorical encoding implies that every primitive has a distinct activity pattern. Consistent with this, activity visualized in a two-dimensional embedding separated primitives in PMv (**Extended Data Fig. 7**). Quantifying this separation, we found that primitives were distinguishable from each other based on single-trial activity (**Extended Data Fig. 12**).

### Recombination of primitives is reflected in PMv

When subjects recombine primitives into multi-stroke sequences to draw characters (**Fig. 2k-p**), does this involve reused primitive-representing states in PMv? We compared activity when a primitive was used in the Single-shape task with activity when the same primitive was used in the Character task. Because Character tasks involve multiple strokes, to facilitate comparison with Single-shape tasks, we focused analysis on the time window immediately preceding stroke onset (rather than the entire planning epoch). We observed that each primitive’s PMv population activity (inner rows in **Fig. 6a**, trajectories in **Fig. 6b**) was similar across task types (columns in **Fig. 6a, b**). In contrast, in preSMA, an area known to encode sequence-related information^57^, activity differed between task types (**Fig. 6c, d**). Quantification showed that PMv encodes primitives similarly across task types. This is evident in the off-diagonal streak in the matrix of pairwise neural distances between primitive-task conditions (**Fig. 6e**), and in the finding that neural distance between primitives controlling for task (“primitive encoding”) was high, while the distance between task types controlling for primitive (“task-type encoding”) was low (**Fig. 6f**). In contrast, preSMA activity differed between Single-shape or Character tasks, which is evident in block structure in **Fig. 6e** and in relatively high task-type encoding compared to primitive encoding (**Fig. 6f**). Across all areas, PMv most consistently exhibited high primitive encoding and low task-type encoding (**Fig. 6f**).

**Fig. 6.**
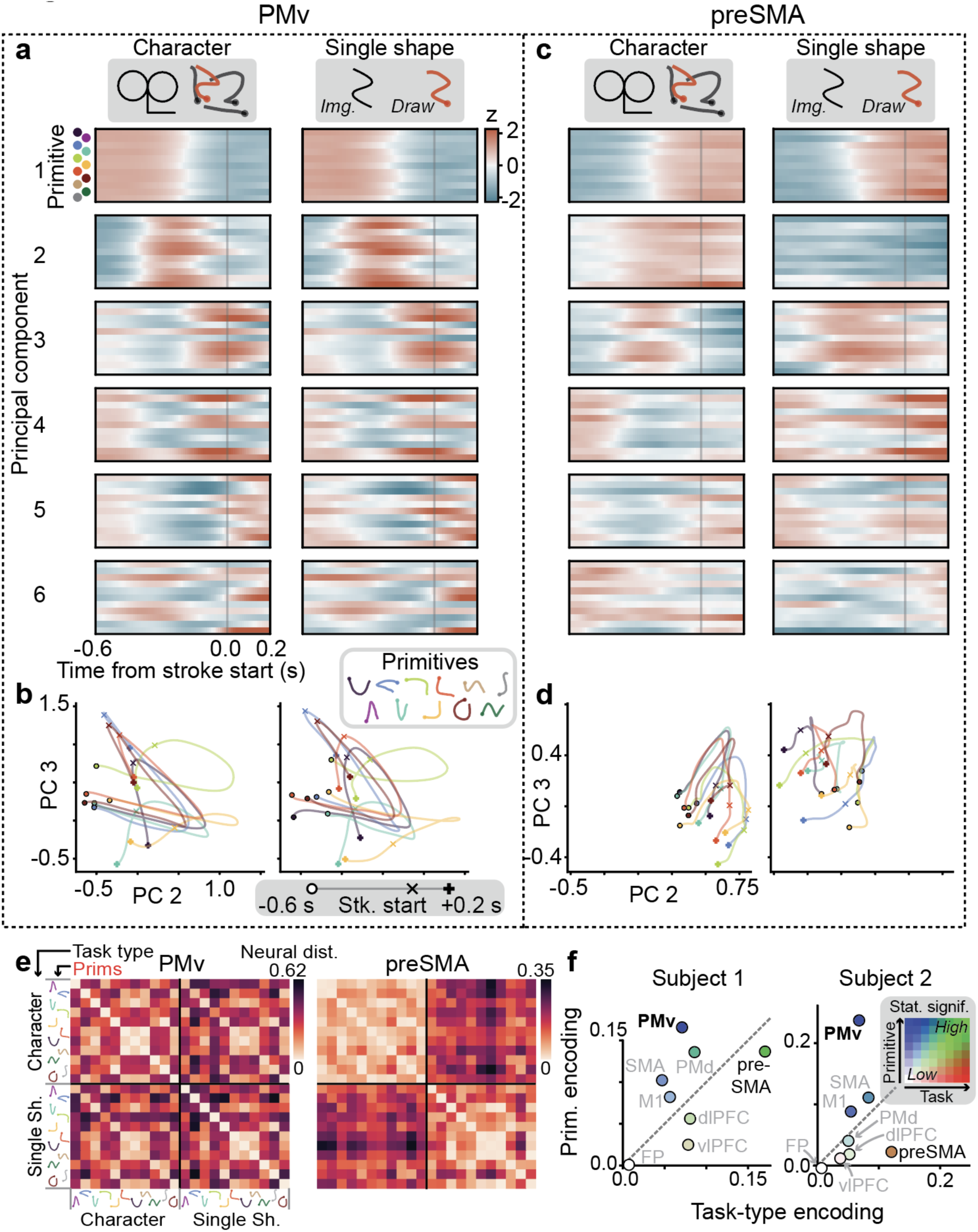
Recombination of primitives into sequences is reflected in PMv. (a) PMv population activity in an example experiment, for its first six principal components (outer rows), split by primitive (inner rows), and task type (columns). Here, activity was aligned to stroke onset and z-scored over the entire window. n (trials) = 4 - 16 (Single-shape), 6 - 72 (Character), in subject 2. Colors index the primitives in panel b. (b) PMv trajectories for principal components 2 and 3, aligned to stroke onset (marked “x”). Gray legend depicts a trajectory. This shows a subset of primitives (7/11) to reduce clutter. (c) Analogous to panel a, but for simultaneously recorded activity in preSMA. (d) Analogous to panel b, but for preSMA. (e) Pairwise neural distance between combinations of primitive and task type, for the experiment in panels a-d, averaged over time (-0.5 to -0.05 s). n (trials) = 4 - 16 (Single-shape), 6 - 72 (Character). (f) Summary of primitive and task-type encoding across areas and sessions. Points show mean encoding scores, with color denoting statistical significance (as in Fig. 4j). Statistical tests were performed on condition pairs pooled across sessions (9 for S1, 9 for S2). We used bootstrapping to balance sample sizes across sessions. Each data point was a unique pair of primitive-task conditions. Primitive encoding (same task type, different primitive) [n = 700 (subject 1), 548 (subject 2)]. Task-type encoding (same primitive, different task type): [n = 82 (S1), 64 (S2)]. For exact effect sizes and p-values, see **Extended Data Fig. 13**.

In the preceding analysis, to account for the fact that Single-shape strokes involve a large initial reaching movement from the start button towards drawing, we included only the first Character stroke, which also involves this reach, to ensure a fair comparison by accounting for neural effects associated with this reach^59,62^. We further expanded this analysis to include all character strokes. We first applied a linear correction to neural activity to account for an additive effect of the initial reach (**Extended Data Fig. 14a,b**). As in the previous analysis (**Fig. 6f**), we found that PMv exhibited high primitive encoding and low task-type encoding (**Extended Data Fig. 14c-e**). These findings indicate that representations of primitives in PMv are reused in recombined sequences.

### PMv activity dissociates from visual and motor parameters

That the effects for motor invariance (**Fig. 4**), categorical structure (**Fig. 5**), and recombination (**Fig. 6**) were strongest in PMv is evidence for a representation of stroke symbols in PMv. We wished to further characterize this representation. PMv can respond to visual object stimuli^34^ and encode movement kinematics^63^, raising the question of how visual and motor parameters contribute to PMv in the current task.

First, we considered whether PMv is driven by the shape in view. We used a task that dissociates vision from action (Multi-shape task). Images consist of multiple separated shapes, with no imposed rule for drawing order. During planning, gaze consisted of saccades and fixations onto different shapes (**Extended Data Fig. 15a-c**). Collecting planning-related fixation events across trials with different first-drawn primitives dissociated the visually-fixated shape from the primitive the subject planned to draw first. We found that PMv strongly encoded the planned primitive regardless of the visually-fixated shape (**Extended Data Fig. 15d-f**), and in a manner largely unaffected by ongoing saccades (**Extended Data Fig. 15d)**. Second, we tested whether PMv encodes stroke kinematics in a manner that generalizes across primitives. We found that this was not the case. A linear encoding model (mapping from stroke velocity to neural activity) trained on one subset of primitives did not explain any variance in PMv activity for held-out primitives (**Extended Data Fig. 16**). In contrast, kinematics accounted for a significant fraction of variance in M1, consistent with its role in motor control. These results show that PMv activity in this task exhibits a form of abstraction in which it is independent from immediate visual input and motor output.

## Discussion

### Identification of a neural substrate of action symbols

We tested for three essential properties of symbols—invariance, categorical structure, and recombination—in behavior and neural activity. In behavior, monkeys traced figures by recombining stroke primitives into new sequences (**Fig. 2k-p**), and these primitives exhibited motor invariance (**Fig. 2a-e**) and categorical structure (**Fig. 2f-j**). In motor, premotor, and prefrontal population activity (**Fig. 3**), we found evidence for these three properties, which, strikingly, were all strongest in a single area, ventral premotor cortex (PMv; **Figs. 4-6**). These behavioral and neural findings reveal a localized representation of action symbols in PMv.

Finding action symbols in PMv was unexpected given prior evidence for encoding of abstract cognitive variables in other areas, especially PFC^8,32,36–38^ and medial temporal lobe^29,31,32,40^. This may reflect a fundamental difference in the role of motor behavior between our study and these prior studies. In our task, complex movement details are relevant for task success, whereas commonly-studied cognitive tasks usually involve simple movements whose main purpose is for reporting a choice (e.g., button press). This suggests that PMv’s critical role is in encoding abstract representations of skilled movements. A privileged role for PMv in motor abstraction is supported by findings in humans, including that (i) PMv lesions are associated with disrupted action conceptual knowledge^64^; (ii) PMv activation is associated with action-related perception^65,66^, imitation^66,67^, verbalization^68^ and imagination^65,66^; and (iii) PMv dysfunction is implicated in apraxia, a disorder of complex motor skills such as drawing^69^. These observations may reflect processes involving action symbols in PMv. More generally, our findings highlight a kind of abstraction that is relatively understudied in systems neuroscience (motor abstraction) and establish a starting point for studying its neural basis (action symbols in PMv).

Our finding in monkeys may provide general insights into the neural basis of symbols. Certain abstract properties—such as reversible symbol reference^70^, higher-order relations^8,71^ and recursive syntax^7,8^—are thought to characterize symbol systems in human language, mathematics, and formal reasoning, and have not been readily apparent in animal behavior^7,8,70,71^, suggesting they may be specific to humans. At the same time, various animal behaviors appear to reflect (possibly less abstract kinds of) symbols, and invariant neural representations have been identified in animals performing cognitive tasks (see Introduction). Finding symbolic representations in macaque motor behavior raises the possibility that symbolic operations exist across species and may reflect shared mechanisms for generating and recombining discrete, invariant representations. Our study’s criteria of invariance, categorical structure, and recombination may be useful in future studies, including in reassessing findings of invariant activity in cognitive tasks as symbols.

### Action symbols and abstraction in PMv

Several lines of evidence indicate this action symbol representation is not directly driven by visual features or motor parameters. First, we found that PMv encoded the planned primitive in a manner dissociated from visual features, when (i) the shape in view was dissociated from the planned primitive (**Extended Data Fig. 15**); (ii) an ambiguous shape was drawn as different primitives (**Fig. 5f**); and (iii) different images were drawn with the same primitive (**Fig. 6f**, Character vs. Single-shape tasks). Second, independence from motor parameters is indicated by three properties of primitive-encoding activity in PMv: (i) presence during planning, temporally dissociated from drawing (by > 1 sec; e.g., **Fig. 5h**), (ii) invariance to location and size (**Fig. 4, Extended Data Fig. 9, 10**), and (iii) lack of encoding of generic motor kinematics (**Extended Data Fig. 16**). These properties in PMv contrast the motor-related activity in M1: (i) lack of primitive-encoding during planning (**Fig. 4j**); (ii) lack of location-invariance during initial reach (**Extended Data Fig. 9**); and (iii) encoding of generic motor kinematics (**Extended Data Fig. 16**). Thus, PMv encodes action symbols in an abstract manner.

Prior evidence for motor invariance in PMv may reflect action symbols. Recordings have revealed a diversity of invariant firing patterns, especially during grasping and object manipulation^34^, including invariance to effector^72^ and muscle activity patterns^63,73^. PMv also contains so-called “mirror neurons” that fire similarly whether one observes or performs a given action^74,75^. These abstract properties are proposed to support a variety of functions, including visuo-motor transformation^34,76,77^, action understanding^74^, imitation^74^, and, perhaps most related to action symbols, an encoding of a repertoire of action types^34,74,76,78^. There are differences between these proposals and the action symbols framework, both conceptually—action symbols place stronger emphasis on categorical structure and recombination—and experimentally—we tested categorical structure (in a systematic manner not previously done) and novel recombination (extending previous studies, which tested only a few well-practiced sequences^79^). Also essential to revealing specificity to PMv was that we recorded from more areas than in prior studies. Our findings raise the possibility that PMv’s involvement in diverse motor behaviors—including speech^80^—may reflect diverse kinds of action symbols.

PMv is at a unique intersection of motor, cognitive, and sensory circuits. Same as other premotor areas (including PMd and SMA), PMv is directly connected with M1 and spinal cord^81^, affording direct control over motor output. In *contrast* to other premotor areas, PMv is strongly interconnected with prefrontal cortex (especially vlPFC)^82^, affording direct access to cognitive processes. PMv is also interconnected with preSMA, an area involved in action sequencing^82^. PMv, vlPFC and preSMA are components of the macaque homologue of the “multiple-demand” system^83^, areas implicated in abstract reasoning. PMv receives input from the two cortical visual pathways: (i) spatial- and action-related “dorsal stream” signals via inferior parietal cortex^84^, and (ii) shape- and object-related “ventral stream” signals via vlPFC^85^. Although ventral stream signals can encode image shape parts^86^ with location and size invariance^87^, and categorical specificity^86,88^, these inputs are unlikely to fully account for these properties in PMv, as we showed that PMv represents the planned action rather than visual input and that vlPFC did not exhibit these properties. In addition to these inputs, processes within PMv may be critical, including winner-take-all competition, which may account for slow activity dynamics for ambiguous images (**Fig. 5h**)^60^ and relate to PMv’s involvement in decision-making^89,90^. Future studies may elucidate how diverse inputs interact with local mechanisms in PMv to produce action symbol representations.

### Motor behavior as a model system for compositionality

Our study introduces a task for studying compositional generalization. We highlight three methodological features advantageous for studying compositional generalization. First, the task required subjects to decide their own actions (no cues instructing movement details) and disincentivized memorization (highly varied, including novel, images). These aspects, coupled with an inductive bias for knowledge compression^7,11,12^, incentivized the learning of generalizable motor concepts (action symbols), and also contrast with prior drawing-like macaque studies, which involved direct instruction using a moving cue that the monkey’s movements had to track^91,92^ or low stimulus variability (two to four highly practiced images)^93^. Second, images were often ambiguous, which was useful for probing the nature of each subject’s prior knowledge. Third, stimulus design was flexible, allowing us to present images that vary parametrically and continuously to test behavior diagnostic of invariance, categorical structure, and recombination. Together, these task features (free choice, stimulus ambiguity, and parametric stimulus design) relate to a critical aspect of complex naturalistic behavior—the need to generalize to situations where behavior is not uniquely determined by sensory inputs and specific prior experiences, but instead depends on internal decision-making computations involving multiple interacting processes, including perception, inference, and efficient planning, all constrained by structured prior knowledge, including of symbols and how they can be flexibly recombined. Thus, we established a task paradigm that allowed us to identify action symbols, and which, in future work, allows studying how multi-area neural processes underlie the internal computations that map from complex sensory input to structured symbolic behavior.

This drawing task complements tasks in prior neural studies of motor behavior, which, to our knowledge, have not directly addressed compositional generalization. They have mainly studied behaviors that are automatic, instructed, working-memory-guided, or minimally-restrained (non-mutually-exclusive classes that expand those in Mizes et al.^94^). Automatic behaviors involve highly-practiced stereotyped sequences^49,52,94–97^. Instructed behaviors are guided by cues that trigger specific actions, either directly (e.g., cue location instructing reach location)^59,62,94,98^ or through learned rules^20,99^. Working-memory-guided behaviors are reproduced from short-term memory^36,57,94,95^. Minimally-restrained behaviors are spontaneously produced, often in a naturalistic setting^49,50,78^. Previous drawing-like tasks in monkeys can be classified as automatic^93^, instructed^91,92^, or minimally-restrained^92^. Thus, our drawing task opens a new direction: compositional generalization.

Behavioral studies suggest that continuous movements can be decomposed into reused segments called “motor primitives”^100,101^. Compared to action symbols, motor primitives are at a lower level of abstraction, often defined as muscle co-activity patterns (“synergies”)^100,101^. An open question is whether motor primitives are encoded in the brain (see evidence in M1^54,62,98,102,103^), or instead are behavioral byproducts of biomechanical constraints, task low-dimensionality, or spinal circuit properties^101^. Related studies have identified reused segments (or “syllables”) in minimally-restrained behaviors, including in birdsong^23,49,50^, which in some cases are idiosyncratic to individuals^104^, similar to stroke primitives in our study. Future studies may address whether motor primitives and syllables reflect action symbols.

### Bridging symbols and neural computation

Our findings may help bridge two paradigms that have dominated the modeling of cognition: one based on symbols and rules^2–8^ and the other on neural network (“connectionist”) architectures and dynamical systems^24,25,27,28^. Explaining cognition may depend on their unification, possibly by implementing symbolic programs^4,5,7,12,37,105^ in neural representations and dynamics^15,26,37,44,105–107^, and incorporating insights from cognitive-task-optimized network models^25,28,108^. Building on our results, future studies may test for activity consistent with symbolic programs in PMv and connected areas, including PFC and preSMA, areas known to encode sequential context [in prior studies^36,37,57,96^ and the current study (preSMA, **Fig. 6**)], and in hippocampal circuits implicated in generating compositional representations^6,41,42,109^.

## Methods

### Subjects and surgical procedures

Data were acquired from two adult male macaques (*Macaca mulatta*, average weights 17 kg (S1) and 10 kg (S2), average ages 9 years (S1) and 7 years (S2)). All animal procedures complied with the NIH Guide for Care and Use of Laboratory Animals and were approved by the Institutional Animal Care and Use Committee of the Rockefeller University (protocol 24066-H).

After undergoing initial task training in their home cages, subjects underwent two surgeries, the first to implant an acrylic head implant with a headpost, and the second to implant electrode arrays. Both surgeries followed standard protocol, including for anesthetic, aseptic, and postoperative treatment. In the first surgery, a custom-designed MR-compatible Ultem headpost was implanted, surrounded by a bone cement cranial implant, or “headcap” (Metabond, Parkell and Palacos, Heraeus), which was secured to the skull using MR-compatible ceramic screws (Rogue Research). After a six month interval, to allow bone to grow around the screws and for the subject to acclimate to performing the task during head fixation via the headpost, we performed a second surgery to implant 16 floating microelectrode arrays (32-channel FMA, Microprobes), following standard procedures^110^. Briefly, after performing a craniotomy and durotomy over the target area, arrays were inserted one by one stereotactically, held at the end of a stereotaxic arm with a vacuum suction attachment (Microprobes). Using vacuum suction allowed us to release the arrays, after insertion, with minimal mechanical perturbation by turning off the suction. After all arrays had been implanted, the dura mater was loosely sutured and covered with DuraGen (Integra LifeSciences). The craniotomy was closed with bone cement.

We used standard density arrays (1.8 mm x 4 mm) for all areas, except SMA and preSMA, for which we used four high density arrays (1.6 mm x 2.95 mm). Four additional electrodes on each array served as reference and ground. Two arrays were targeted to each of multiple areas of frontal cortex, with locations identified stereotactically, and planned using brain surface reconstructions derived from anatomical MRI scans. Locations were selected based on their published functional and anatomical properties (see below), anatomical sulcal landmarks, and a standard macaque brain atlas^111^. During surgery, locations were further adjusted based on cortical landmarks, and to avoid visible blood vessels. Arrays were implanted in the right hemisphere (contralateral to the arm used for drawing).

Array locations are depicted in **Fig. 3b** (confirmed with intraoperative photographs). For M1, we targeted hand and arm representations (F1), directly medial to the bend of the central sulcus (which corresponds roughly to the intersection of the central sulcus and the arcuate spur if the latter were extended caudally), based on retrograde labeling from spinal cord and microstimulation of M1^81^ and M1 recordings^112^. For PMd, we placed both arrays lateral to the precentral dimple, with one (more caudal) array directly medial to the arcuate spur (the arm representation^81,112,113^, F2), and the other was placed more rostrally (straddling F2 and F7). For PMv, we targeted areas caudal to the inferior arm of the arcuate sulcus (F5), which are associated with hand movements based on retrograde labeling from spinal cord^81^ and M1^114^, microstimulation^115^ and functional studies^77,89,116^. These areas contain neurons interconnected with PFC^114^. For SMA (F3) and preSMA (F6), we targeted the medial wall of the hemisphere, with the boundary between SMA and preSMA defined as the anterior-posterior location of the genu of the arcuate sulcus, consistent with prior studies finding significant differences across this boundary in anatomical connectivity (e.g., direct spinal projections in SMA but not preSMA^117^) and function^57,118^. SMA arrays were largely in the arm representation^117^. For dlPFC, we targeted the region immediately dorsal to the principal sulcus (46d), following prior studies of action sequencing^36,37,119^ and other cognitive functions^120^. For vlPFC, we targeted the inferior convexity ventral to the principal sulcus, with one (more rostral) array directly ventral to the principal sulcus (46v) and the other rostral to the inferior arm of the arcuate sulcus (45A/B), based on evidence for encoding of abstract concepts in regions broadly spanning these two locations^38,121,122^, including a possible heightened role (compared to dlPFC) in encoding abstract concepts in a manner invariant to temporal or spatial parameters^122–124^. For FP, we targeted a rostral location similar to prior recording and imaging studies (one array fully in area 10, the other straddling 9 and 10)^125,126^. In general, array locations targeted the cortical convexity immediately next to sulci, instead of within the banks, in order to allow shorter insertion depths that minimize the risk of missing the target. The exceptions were SMA and preSMA in the medial wall, for which this was not possible. To avoid damaging the superior sagittal sinus, we positioned the arrays laterally (2 mm from midline) and slanted the electrodes medially (**Extended Data Fig. 6**).

The lengths of each electrode were custom-designed to target half-way through the gray matter, and to vary substantially across the array, to maximize sampling of the cortical depth. Electrode lengths spanned 1.5 - 3.5 mm (M1), 1.5 - 3.1 mm (PMd, PMv), 2.8 - 5.8 mm (SMA, preSMA), 1.5 - 2.5 mm (dlPFC, vlPFC), and 1.5 - 2.6 mm (FP) for subject 1, and 1.7 - 3.75 (M1), 1.5 - 3.3 mm (PMv), 1.5 - 3.1 mm (PMd), 2.65 - 5.95 mm (SMA, preSMA), 1.75 - 3.15 mm (dlPFC), 1.35 - 3.2 mm (vlPFC), and 1.6 - 2.9 mm (FP) for subject 2. Reference electrodes were longer (6 mm) to anchor the arrays. 28 electrodes were Pt/Ir (0.5 MΩ) and 4 were Ir (10 kΩ) (for microstimulation). Array connectors (Omnetics A79022) were housed in custom-made Ultem pedestals (Crist), which were secured with bone cement onto the cranial implant. Four pedestals were used per subject, holding 5, 5, 4, and 2 connectors each.

### Behavioral task

#### Task overview

Subjects were seated comfortably in the dark with their head restrained by fixing the headpost to the chair. They faced a touchscreen (Elo 1590L 15” E334335, PCAP, 768 x 1024 pixels, refresh rate 60 Hz, with matte screen protector to reduce finger friction) that presented images and was drawn on. The touchscreen location was optimized to allow each subject to easily draw at all relevant locations on the screen (23 to 26 cm away; see diagram in **Extended Data Fig. 1**). Both subjects decided on their own over the course of learning to perform the task with the left hand. The chairs were designed to minimize movements of the torso and legs (by using a loosely restricting “belly plate”), and of the non-drawing arm (by restricting movement to within the chair). Gravity-delivered reward (water-juice mixture) was controlled by the opening and closing of a solenoid pinch valve (Cole-Parmer, 1/8" ID). Subjects were water-regulated, with careful monitoring that consumption met the minimum requirement per day (and typically exceeded it), and weight was closely monitored to ensure good health. The task was controlled with custom-written software, using the MonkeyLogic behavioral control and data acquisition MATLAB package^127^. (PC: Windows 10 Pro, Intel Core i7-4790K, 32GB RAM; DAQ: National Instruments PCIe-6343). All stimuli (images of line figures defined as point sets) were also generated with custom-written MATLAB code. Images were presented in a “workspace” area on the screen (16.6 cm x 16.9 cm, corresponding to approximately 37° by 38° visual angle). Shape components in images were on average 4.0 cm (9°) (taking the maximum of width and height).

Each neural recording session consisted of a day’s recording (2 - 3.5 hours). We collected 5-20 trials per condition (i.e., each unique image for **Figs. 4, 5** and Single-shape tasks in **Fig. 6**; each primitive stroke for the Character task in **Fig. 6**). All trials were shuffled across all conditions within the session, and presented in a randomly interleaved fashion, except for one case, the experiment in **Fig. 6**, in which Character and Single-shape tasks were switched in blocks.

#### Early Training

Before surgery, naive subjects underwent initial training on core task components (i.e., to trace images accurately on a touchscreen, using a sequence of discrete strokes). Early training took place in the home cage, using custom-built rigs that attached to an opening in the cage, using the same hardware and software described above, except for a different computer (Lenovo IdeaPad 14" laptop, Windows 10, AMD Ryzen 5 3500U, 8GB RAM) and DAQ (National Instruments USB-6001). This initial training progressed through the following stages: *(1) Touch circle*. Subjects were rewarded for touching a circle anywhere within its bounds. The circle started large, filling the entire screen, and shrank over trials to enforce more accurate touches. *(2) Touch with a single finger*. We shrank the circle until it was so small that it could only be touched with a single finger. The trial aborted if the subject touched outside the circle, or with multiple fingers simultaneously. *(3) Hold still*. Subjects were rewarded for keeping their fingertip still on a dot, with the duration of this hold increasing across trials (up to a few seconds). *(4) Track moving dot.* Subjects had to track the dot with their finger as it moved (a lag between dot and finger was allowed). *(5) Trace a line.* We increased the speed of the moving dot over trials, until eventually the dot moved so fast that the line it traced appeared immediately. We then positioned the line at locations far from the hold position to train the subject to raise its finger from the hold position and trace lines at arbitrary locations, angles, and lengths. *(6) Trace single shape.* We presented shapes of increasing difficulty (gradually “morphing” across trials from a straight line), including arcs, “L”-shapes, and squiggles, and circles. Across these stages, the shapes were presented at random locations. We did not enforce any particular tracing strategy for each shape (e.g., which endpoint to start at), allowing subjects to choose on their own. *(7) Trace multiple shapes.* We presented images composed of multiple disconnected shapes. This trained the subjects to understand that they should use multiple strokes to trace multiple shapes. At this point, the subject understood the basic structure of the task—to trace shapes, using multiple strokes if needed. The progression across these stages was not determined by strict quantitative criteria, but instead on a combination of quantitative and qualitative evaluations of how well the subjects understood the task.

After this basic training, subjects practiced various tasks to incentivize the learning of stroke primitives (consistent stroke trajectories for each shape). They practiced Single-shape trials using the set of diverse simple shapes in **Fig. 1e**, varying randomly in shape and location across trials. We chose this set of shapes in order to cover a variety of trajectory profiles (by varying rotation, the number of direction changes, and whether shapes were curved or linear), and yet to still be simple enough to draw with one stroke and to be able to combine multiple (two to six shapes) shapes into single character images. We did not constrain subjects to learn specific stroke trajectories for each primitive; therefore, differences between primitives reflected each subject’s own decision for how to draw each shape. We also note that our study did not depend on using an “optimal” set of shapes, but instead depended on subjects learning primitives associated with the basic shapes, and then demonstrating behavioral generalization using these primitives, as we found that they did. For subject 2, the four “S” shapes and four “arc” shapes were also sometimes presented as rectilinear versions. For “S” these resembled “Z”, and for “arc” these resembled squares missing one edge. The subject drew these rectilinear versions using curved strokes similar to those used to draw “S” and “arc” shapes. Therefore, instead of considering these rectilinear versions as distinct primitives, we combined their data with their respective curved versions. On different days, subjects also practiced Multi-shape and Character tasks. Below, we describe the trial structure, followed by details for Single-shape, Multi-shape, and Character tasks.

#### Trial structure

As depicted in **Fig. 1c**, trials began when the subject pressed and held a finger fixation button (blue square) at the bottom of the screen (gray background; note that in this article “button” always means a virtual button; diagram of screen in **Extended Data Fig. 1c**). After a random delay (uniform, 0.2 s window, earliest [0.4, 0.6] s and latest [0.8, 1.0] s across experiments for subject 1 and [0.8, 1.1] s for subject 2) the image appeared (figure colored dark gray). After a random delay (uniform, ranging from [0.6, 1.0] s to [1.2, 1.6] s across experiments for subject 1, and [1.1, 1.5] s to [1.8, 2.4] s for subject 2), a “go” cue (400 Hz tone and image blank for 300 ms) was presented. This delay between image presentation and go cue we call the “planning” epoch. The subject had to keep its finger still on the fixation button during the planning epoch. After the onset of the go cue, the subject was free to raise its finger, move its hand towards the image, and start drawing. During drawing, the image stayed visible, and the finger left a trail of black “ink” on the screen. Immediately after the finger was raised from the fixation button after the go cue, the “fixation button” disappeared and a “done button” (green square) appeared at its location and stayed on. The drawing epoch ended when the subject signaled it had completed the drawing, by pressing the “done button” (no time limit was imposed). This was followed by a delay (uniformly sampled, 0.65 to 1 s), followed by performance feedback. Feedback spanned four modalities, each signalling performance: (i) screen color, (ii) sound cue, (iii) duration of a delay before getting reward, and (iv) reward magnitude. First, screen color and sound were signaled, followed by the pre-reward delay and then the reward. How performance was scored and converted to feedback is described below. In addition to providing this feedback at the end of the trial, we also provided feedback online by immediately aborting a trial if subjects made serious errors, including (i) if they touched a position far from any image points, and, (ii) for Single-shape and Multi-shape trials, if they used more than one stroke per shape in the image. These “online abort” modes were turned off for trials testing novel characters.

Screen image changes (including image presentation and trial events) were recorded using photodiodes (Adafruit Light Sensor ALS-PT19) and sounds were recorded using an electret microphone (Adafruit Maxim MAX4466, 20-20KHz). We performed eye tracking (ISCAN), but did not enforce eye fixation.

#### Scoring behavioral performance

Behavior was scored by aggregating multiple metrics, or “factors”. There were three classes of factors. The primary class measured *image similarity*, or the similarity of the final drawing to the target image (ignoring its temporal trajectory). This factor had the greatest influence on the final aggregate score. Additionally, we computed factors reflecting *behavioral efficiency*, and, in some cases, factors that were *task-specific*. These scores were computed on behavioral data, which was represented as a sequence of touched points (x-y coordinates) with gaps between strokes, and image data, represented as a set of x-y coordinates. First, we describe the factors, and then how they were aggregated into a single score.

##### Image similarity

This included two factors: drawing-image overlap and Hausdorff distance. Drawing-image overlap was computed as the fraction of the image points that were “touched” (within a margin of error) by at least one of the drawn points. A subset of the image points were weighted more heavily because they captured unique features of the shape (e.g., the corners and endpoints of an “L” shape). Hausdorff distance is a distance metric between the set of drawn points and the set of image points (see definition below).

##### Behavioral efficiency

To incentivize efficiency, we included a factor comparing the cumulative distance traveled in the drawing (i.e., the amount of “ink”) to the cumulative distance of the edges of the figure in the image, with its value negatively proportional to the excess of drawn ink over image ink.

##### Task-specific factors

During practice trials for the Character task (see “Task types”), we also included factors capturing the extent to which drawn strokes matched the shapes used in the image figure. This included two factors: one proportional to the similarity of the number of strokes and the number of image shapes, and the other proportional to the spatial alignment of the drawn strokes to the image shapes. Importantly, these factors were included only for practice images, not for novel test images.

The final score aggregated the image similarity, behavioral efficiency, and task-specific factors, with more weight on image similarity factors. We first rescaled the factors linearly between 0 and 1 (where 1 means good performance), with the dynamic range set by a lower and upper bound on the factor values. These bounds were adaptively updated on every trial based on the distribution of factor values in the last 50 trials (lower bound at 1st percentile and upper bound at 53rd percentile), which ensured that the dynamic range of feedback matched the dynamic range of behavioral performance from recent history. We then weighted each of those factors to tune their relative contributions (using hand-tuned weights for each experiment; generally highest for image similarity), and computed the final scalar score (range 0 to 1) by taking the single factor that was worst after weighting.

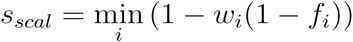

where *i* indexes the factors, *w_i_* are the weights (between 0 and 1) and *f*_1_ are the factor values (rescaled between 0 and 1). We also gave each trial a categorical score:

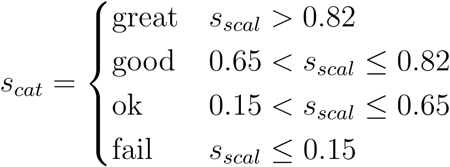

The scalar and categorical scores were used to determine feedback across four different modalities. The meaning of screen color and sound were learned by the subject, whereas delay and reward had intrinsic value:

*(1) Screen color.* A linear mapping between two colors, such that a score of 0 was mapped to red (RBG: [1, 0.2, 0]) and a score of 1 was mapped to green ([0.2, 1, 0.2]).
*(2) Sound cue.* A sound determined by *s_cat_*: if *great*, then three pulses (1300 Hz, 0.16 s on and off); if *good*, then a single pulse (1000 Hz, 0.4 s); if *ok*, then no sound; if *fail*, then a single pulse (120 Hz, 0.27 s).
*(3) Delay until reward.* A nonlinear mapping from score to delay before reward. We first applied a linear mapping to the scalar score, such that a score of 0 was mapped to a long delay (5 sec + random uniform jitter, [0, 2.5] s) and a score of 1 was mapped to 0 s delay. Further, if *s_cat_* was *great*, *good*, or *ok*, then this delay was reduced by multiplying by 0.65.
*(4) Reward.* The open duration of the solenoid gating the juice line was defined as

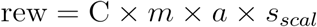

where C is a constant in dimensions of time (0.15 - 0.6 s, set manually depending on the difficulty of the task); *m* is a multiplier that gives a bonus for good performance and further penalizes bad performance, depending on the value of *s_cat_*: *great* (1.3), *good* (1.0), *ok* (0.8), *fail* (0); and *a* is a random variable sampled from the uniform distribution *a* ~ 0.75 + 0.5 × *U*(0,1), and *s_scal_* defined as above. On average, including failed trials, subjects received ∼0.35 ml reward per trial. The order in which these four feedback signals were delivered is described above (“Trial structure”).

#### Task types

##### Single-shape task

The Single-shape task presented one of the practiced simple shapes, or, in the “categories” experiments, sometimes a morphed shape. Subjects were allowed only a single stroke (triggering online abort if >1). In four Single-shape sessions for subject 1, the ending of the drawing epoch was triggered by completion of the stroke (i.e., on finger raise), and not on the subject pressing a “done” button, as was the case in all other sessions and experiments.

To test for motor invariance (**Figs. 2a-e, 4**), we presented images of practiced shapes, varying across trials in location, size, or both. For location variation, images spanned a distance in the x- and y-dimensions (measuring from shape centers) of 321 pixels (9.6 cm), which is 2.38 times the average size of shapes (135 pixels, 4.0 cm, maximum across width and height). For size variation, the maximum size was 2.5 times larger than the smallest (in diameter), except for two experiments for subject 1, in which the ratio was 2.0. The location and size variation in **Fig. 2b, c** is representative of all experiments.

This variation in size and location was chosen based on parameters used in prior studies of M1 and electromyographic activity (EMG) of muscles controlling the arm during reaching tasks in macaques. For reaches performed along the coronal plane around 25 cm from the subject, similar to the geometry of the touchscreen relative to subjects in our study, M1 and EMG activity are significantly affected by translating the starting and ending locations of reaches by 10 cm, similar to the location variation in our study (9.6 cm)^128^, and by varying the scale of the reach by 2 fold (7 cm to 12 cm), similar to our study’s variation in shape 2 to 2.5 fold^129^.

To test for categorical structure (**Figs. 2f-j, 5**), we constructed morph sets (subject 1: n = 7 morph sets across 3 sessions; subject 2: n = 13 morph sets across 4 sessions), each consisting of two practiced shapes and four to five “morphed” versions of those shapes, which were constructed by linearly interpolating between the two shapes along one image parameter, such as the extent of closure of the top of the U (**Fig. 2f, g**). Across morph sets, we varied different image parameters (see examples in **Extended Data Fig. 3**).

##### Multi-shape task

Each image was composed of two to four shapes positioned at random, non-overlapping, locations spanning the space of the screen (four corners and center). Subjects were allowed to draw the shapes in any order, and to use any trajectory within the shapes, but were constrained to use one stroke per shape and to not trace in the gaps between shapes. In the Results of this manuscript, we present results averaged across two sessions, one from each subject (**Extended Data Fig. 4**). On each trial, an image was constructed by sampling a shape randomly without replacement. This led to n = 531 (S1) and n = 278 (S2) unique images.

##### Character task

Each image was generated by connecting two to six simple shapes into a single character figure. This was done by sampling characters from a generative model. A character with N shapes was defined by randomly sampling N shapes and N-1 relations, where each relation *i* defines the location of the attachment points on shapes *i* and *i* + 1; these attachment points, in turn, define how these shapes will connect to each other. This is similar to a published generative model for handwritten characters^11^. Generated characters were only kept if there was minimal crossing of shapes over each other.

For experiments testing behavioral generalization to novel characters (**Fig. 2k-p**), we mixed practiced and novel characters (practiced characters: n = 189 (mean, range 22 to 491) per day; novel characters: 48 (mean, range 0 to 155) per day). For analyses, we label as “novel” only the very first trial for a given character. Because of random sampling in generating characters, it would in principle be possible that characters generated on different days are in fact identical; to avoid this, we ensured that all characters we called “novel” indeed did not match any images the subject previously encountered. We did this by ensuring that each novel character was different from every previously encountered character across all days (quantified with the Hausdorff distance).

For neural experiments comparing Single-shape and Character tasks (**Fig. 6**), we analyzed the sessions for which we collected data from both the Single-shape and Character tasks [subject 1: n = 10 sessions, median N matching primitives between Single-shape and Character tasks = 9 (range 5 - 12); subject 2: n = 9 sessions, median N matching primitives = 10 (range 2 - 14)]. We switched between Single-shape and Character tasks using a block design (2 - 5 blocks each), except one session for subject 2, in which they were randomly interleaved across trials.

### Behavioral data analysis

#### Preprocessing of touchscreen data

Touchscreen data were represented as time series of (x, y) coordinates in units of pixels (conversion: 33.6 pixels/cm) and sampled at 60 Hz, which we upsampled to 500 Hz (performed automatically in MonkeyLogic to align all behavioral signals, including trial event markers and eye tracking), and low-pass filtered below the highest frequency in the drawing movements (15 Hz). Strokes were segmented based on the time the finger first touched the screen (stroke onset) and the last time before raising off the screen (stroke offset) with 500 Hz resolution.

For some analyses—in **Fig. 1f** and as input to the “trajectory distance” below—we further computed stroke instantaneous velocity and speed in the following steps. Starting from extracted strokes, we low-pass filtered the data (12.5 Hz), and downsampled to 25 Hz. We then used the five-point stencil method to compute a finite difference approximation of the derivative:

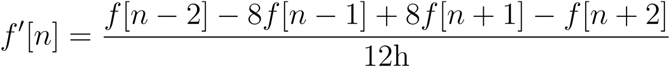

where *f*[*n*] is a discrete time series (i.e., the x- or y-coordinates) indexed by integer *n*, and h is the sampling period in seconds. This differentiation was performed separately for the x and y coordinates. The resulting velocity time series was upsampled to the original 500 Hz sampling rate with a cubic spline. Speed was computed as the norm of the (x, y) velocity at each timepoint.

#### Computing “trajectory distance”

To quantify the similarity between two strokes, in a way that compares their spatio-temporal trajectories, while ignoring their relative size (or scale) and location (on the screen), we devised a “trajectory distance” metric. This metric is a scalar dissimilarity score, based on the dynamic time warping distance between two strokes represented as velocity time series *v*_1_ and *v*_2_. To compute trajectory distance between two strokes, we (1) spatially rescaled each stroke (while maintaining its x-y aspect ratio), to make the diagonal of its bounding box unit length 1. (2) We then linearly interpolated each stroke to the same number of points (70) to allow direct point-by-point comparison between strokes. This was done spatially by interpolating based on the fraction of cumulative distance traveled (so that the distances between successive points were the same value over the entire stroke), in order to capture the spatio-temporal trajectory, as in a previous study modeling strokes in handwriting^11^. (3) Interpolated trajectories were then converted to velocity time series as above. (4) We then computed the dynamic time warping distance between these velocities *v*_1_ and *v*_2_.:

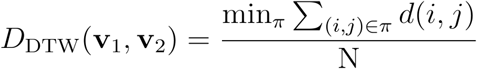

where *i* and *j* index the two velocity trajectories, *N* is the number of points (70), *π* is a set of (*i*,*j*) pairs representing a contiguous path from (0, 0) to (N, N). The local distance metric *d*(*i*,*j*) is the Euclidean distance plus a regularization factor that discourages excessive warping:

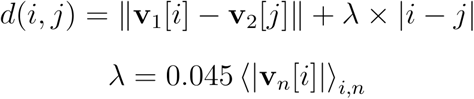

where ⟨·⟩ is the average. For the regularization parameter, *λ*, the purpose of the summation term was to rescale lambda to match the magnitude of velocities. The resulting distance *D_DTW_* was then rescaled to 0 and 1 to return the trajectory distance:

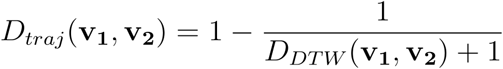

#### Computing “image distance”

To compare the similarity of two images—each a figure represented as a set of (x, y) points, with no temporal or stroke-related information—we used a modified version of the Hausdorff distance, a distance metric that has been commonly used in machine vision for comparing the similarity between two point sets based on their shape attributes^130^. There are, in principle, at least 24 variants of the Hausdorff distance based on possible variations in the formula^130^; we used a variant that is minimally susceptible to outlier points (because it is based on taking means instead of minima and maxima; variant #23 in the referenced study^130^). Image distance was computed as follows: (1) Each image was first centered so that its center of mass becomes (0,0). (2) Image distance was then computed. First, we define the distance between two points, *d*(*a*, *b*), as the Euclidean distance. We also define the distance between a point and a set of points, *d*(*a*, *B*), and the distance from set A to set B, *d*(*A*, *B*), as:

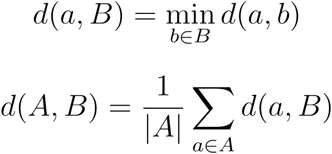

Image distance was then defined as:

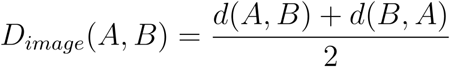

#### Computing “primitive alignment score”

For experiments on categorical structure, we generated a set of images, with each set containing four to five novel images that morph between one primitive (P1) and another primitive (P2): a “morph set”. Across morph sets, different image parameters were morphed (see **Fig. 2g** and **Extended Data Fig. 3**). Each trial presents a single image from one morph set. We sought to quantify the relative similarity between a given trial’s data—either its behavioral, image, or neural data (see below)—and data for the two primitives, P1 and P2, in its morph set. To do so, we devised a “primitive alignment” score defined, for each individual trial, as:

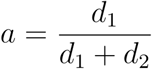

where *d*_1_ is the average of the distances between that trial and each of the P1 trials, and *d*_2_ the average of the distances between that trial and each of the P2 trials. A score closer to 0 implies similarity to P1, and a score close to 1 implies similarity to P2 (note that primitive alignments for P1 and P2 data are not exactly 0 and 1 due to trial-by-trial variation). The particular metric used for these distances depended on the analysis. For images, we used the image distance. For drawings, we used the trajectory distance. For neural activity, we used the Euclidean distance between population vectors. We confirmed that primitive alignment scores for image data varied linearly with morph number (Fig. 2j and Extended Data Fig. 3c), ensuring that any deviation from linearity in behavioral or neural data could not trivially be the consequence of how the score is defined.

#### Classifying strokes from the Character task

To assess whether subjects drew characters by reusing their own stroke primitives, we scored the fraction of character strokes that were high-quality matches to one of the subject’s own primitives, and the fraction that were high-quality matches to the other subject’s primitives. If the fraction of matches to a subject’s own primitives was high, and to the other subject’s primitives was low, then we interpreted this as evidence that subjects recombined their own primitives.

This was performed by assigning each stroke the label of its nearest primitive using the trajectory distance, and then further defining this as a high-quality match only if the trajectory distance was sufficiently low; in particular, if the distance was within the interval bounded by the 2.5th and 97.5th percentiles of the distribution of trajectory distances caused by trial-by-trial variation in behavior.

First, each stroke was assigned its best-matching primitive, *p*\* from a set of primitives (the choice of primitive set—same or different subject—depending on the analysis; see below):

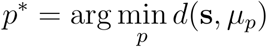

Where *p* indexes the primitives, s is the stroke trajectory, *μ_p_* is the mean stroke for primitive *p* (averaged over trials from Single-shape tasks), and *d*(·, ·) is the trajectory distance.

Second, the quality of the stroke’s assignment to its nearest primitive was scored:

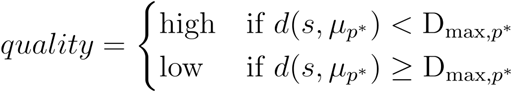

Where D_max,*p*\*_ is an upper-bound on trajectory distances that would be expected from trial-by-trial variation in primitive *p\**. It is the 97.5th percentile of the distribution of trajectory distances from Single-shape trials, determined separately for each primitive, which we consider as a good—if anything, conservative—estimate of trial-by-trial variation, because Single-shape tasks present no ambiguity as to what primitive needs to be drawn.

These steps assigned each stroke a class tuple (*p*, quality*). In summary analyses, we pooled all cases of high-quality matches into a single “high-quality match” class (no matter the assigned primitive), and all low-quality matches into a single “no match” class (**Fig. 2o, p**). In summary analyses testing whether a given subject’s character strokes aligned better with its own primitives vs. the other subject’s primitives (**Fig. 2p**) we performed the above analysis separately for all four combinations of stroke data (2 subjects) x primitives (2 subjects), using only images performed by both subjects.

The control analysis testing primitive reuse using a simulated set of “remixed” primitives was performed in the following manner. We first generated remixed primitive sets by mixing subparts of different primitives. Given two actual primitives we extracted the first half of the first primitive (defined by distance traveled) and the second half of the second primitive and connected them by aligning the offset of the first half to the onset of the second half, smoothing the connection with a sigmoidal weighting function. To sample a remixed primitive set, we first generated a pool of all possible remixed primitives using every possible pair of actual primitives. We then sampled (without replacement) a set of remixed primitives from this pool, keeping only remixed primitives satisfying the following constraints: (i) no self-intersection, and (ii) no excessively sharp turns, detected as curvature at any point along the inner 80% of the stroke exceeding 0.8. Curvature was defined in a standard manner as the inverse of the radius of curvature, (*ẋÿ* – *ẏẍ*)/((*ẋ*^2^ + *ẏ*^2^)^3/2^), where *ẋ* and *ẏ* are the velocity components and *ẍ* and *ÿ* are the acceleration components. Once a candidate set of remixed primitives was sampled, it had to pass further constraints: (i) each actual primitive contributed its first half or second half to a maximum of two remixed primitives; (ii) the trajectory distance between no pair of remixed primitives was lower than the minimum trajectory distance between actual primitives; (iii) the trajectory distance between no pair of remixed primitive and actual primitive was less than the minimum trajectory distance between actual primitives. These constraints ensured that remixed primitives in each set were different from each other and from the actual primitives (this is visually apparent in the primitive sets, **Extended Data Fig. 5**).

We then labeled strokes from character drawings using each remixed primitive set, using the same approach as for actual primitive sets, except the following. The analysis using actual primitives used trajectory distance thresholds determined empirically for each primitive based on drawings in Single-shape trials (see above). Because remixed primitives were not ever drawn, a similar approach could not be used. Instead, thresholds for remixed primitives were assigned from the pool of thresholds for the actual primitives. This was done in a manner meant to increase the match rate (and thus allow a stronger, or more conservative, test that the remixed primitives do not match the character strokes) by assigning the largest (i.e., most lenient) threshold to the “worst”-matching primitive, the second largest to the second worst, and so on. Here, “worse” is defined as having larger average trajectory distance to character strokes.

#### Analysis of kinematic separability of primitives

We tested whether each primitive was decodable from every other primitive based on single-trial kinematics (stroke trajectory) (**Extended Data Fig. 2**). Strokes were first converted from time series (x-and y-position) to 8-dimensional vectors as follows. Similar to the trajectory distance computation (above), strokes were normalized in time (linear interpolation to 50 time points) and space (rescaling each stroke, while maintaining its x-y aspect ratio, to make the diagonal of its bounding box unit length 1). The x- and y- coordinates were concatenated such that each stroke was represented by a 100-dimensional vector, and then reduced to 8 dimensions using PCA. Decoding was performed on these 8D representations using a linear support vector machine classifier (SVC) (LinearSVC, scikit-learn, regularization parameter C set to 0.1), with 10-fold cross-validation. By performing this procedure for every pair of primitives (each time returning a single decoding accuracy score) we populated the matrix in **Extended Data Fig. 2**.

### Neural recordings

Recordings were acquired using a Tucker-Davis Technologies (TDT) system, including headstage (Z-Series 32 Channel Omnetics, LP32CH-Z), amplifier (PZ5M-256), processor (RZ2), and storage (RS4), sampled at 25 kHz (local reference mode), controlled with TDT Synapse software run on a Windows 10 PC (Intel Core i7-3770, 32GB RAM), and saved to disk. Analog and digital task-related signals, including behavioral events (photodiode, audio, and trial event markers) and eye tracking (ISCAN, 125 Hz), were synchronized external triggers recorded by the neural data acquisition system.

### Neural data preprocessing

#### Spike sorting

We extracted for later analysis both isolated single-unit (SU) and multi-unit (MU) spike clusters from the stored broadband signal. MU clusters consisted of threshold crossings that were clearly spikes, but which were not isolatable into distinct SU clusters. We used a three-step approach for extracting and sorting spikes into these clusters, with a first pass using Kilosort^131^ (v2.5) to extract putative spike clusters, a second pass using a custom-written program to label these clusters as either SU, MU, or noise, and a final manual curation step. Note that Kilosort classifies clusters, but we did not use those labels.

For Kilosort, we used the default parameters, except *AUCsplit* (0.90), *Th* ([6, 4]), and *lam* (10), which we optimized using parameter sweeps on data from representative sessions and by manual evaluation of results.

We next refined these cluster labels. For each cluster, we first removed outlier waveforms (any exceeding a 3 x interquartile-range threshold for any of the minima, maxima, or sum-of-squares). Waveforms were then shifted slightly in time (<1 ms) to improve their alignment by peaks (or toughs, for negative-going waveforms). For each cluster, we computed two features. *(1) Signal-to-noise ratio (SNR)*, defined as the ratio of the peak-to-trough difference (of the average spike waveform) divided by standard deviation (averaged across time bins). Before computing SNR, we checked whether the cluster contained both positive- and negative-going waveforms. If so, SNR was computed separately for these two subsets of data and then averaged. *(2) Inter-spike-interval violations (ISIV)*, defined as the fraction of inter-spike intervals less than a refractory period (1.5 ms). Based on these SNR and ISIV, we provisionally classified clusters as SU [if either (SNR > 9.6 and ISIV < 0.05) or (SNR > 6.9 and ISIV < 0.01)], noise (SNR < 3.9), or MU (the remaining clusters).

We then manually curated these clusters. We visualized every cluster to either confirm its label (MU, SU, noise) or to manually re-assign it to a different label (including “artifact”), using a custom-written MATLAB GUI. We also manually checked whether multiple SU clusters on a single channel should be merged into a single SU cluster, if they have high waveform similarity, inversely correlated spike count frequency over the course of the session, or a negative peak close to zero lag in a cross-correlogram of spike times. Finally, for each channel, all MU clusters were merged into a single MU cluster. Combining SU and MU, this yielded the following total number of units per area (mean +/- S.D. across sessions). For subject 1: M1 (59.9 +/- 12.5), PMd (44.1 +/- 6.2), PMv (34.2 +/- 7.3), SMA (63.0 +/- 7.9), preSMA (75.4 +/- 17.7), dlPFC (47.8 +/- 17.2), vlPFC (43.3 +/- 9.9), FP (19.2 +/- 3.8). For subject 2: M1 (40.7 +/- 13.1), PMd (54.7 +/- 5.5), PMv (71.1 +/- 6.6), SMA (53.4 +/- 7.2), preSMA (57.9 +/- 11.1), dlPFC (24.6 +/- 4.8), vlPFC (38.6 +/- 13.7), FP (42.6 +/- 5.0).

#### Converting spike times to firing rates

Single-trial spike trains were converted to firing rate functions by smoothing with a 0.025 s Gaussian kernel (0.01 s slide). We removed units with very low firing rates (if the 80th percentile of their firing rates across all trials and time bins was less than 1 Hz). We also removed units whose firing rates were unstable, by first removing units with high systematic drift in firing rate over the session (*m/u* > 0.2, where *m* is the slope from regressing square-root-transformed firing rate vs. time (in hours), and *u* is the mean firing rate), and then removing units with any large fluctuations in firing rate across the session. For the latter, we excluded units satisfying (*s_max_* - *s_min_*)/*s_mean_* > 1.15 or (*u_max_* - *u_min_*)/*u_mean_* > 0.65, where the session is first split into contiguous 50-trial bins, the across-trial standard deviation in square-root-transformed firing rate is computed within each bin, and then *s_max_*, *s_min_*, and *s_mean_* are defined as the maximum, minimum, and mean standard deviation across bins. *u_max_*, *u_min_*, and *u_mean_* are defined similarly, except using within-bin mean firing rate instead of within-bin standard deviation. We then processed firing rates as follows. We square-root transformed activity to normalize its variance. Following a common approach in analyses of population firing rates^96^, we “soft” z-scored each unit’s activity to ensure that neurons with very different firing rates contributed similarity to population analyses, but with higher-firing-rate neurons still contributing relatively more:

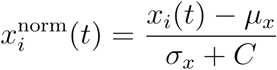

Where *x_i_*(*t*) is firing rate for trial *i* at time bin *t*, *μ_x_* and *σ_x_* are the mean and standard deviation, respectively (across trials and time bins), and *C* is an additive factor to ensure “softness”: *C* = min(**m**) + 3Hz where m is a vector of mean firing rates, one for each unit. All subsequent analyses used this normalized firing rate representation of the data.

#### Time-warping neural activity to a common trial template

For the figure showing the average firing rates over the entire trial (**Fig. 3**), we first time-warped each trial to a common trial template. We defined a set of events that occur across trials as “anchors” (fixation touch, image onset, go cue, finger raise off fixation, stroke onset, stroke offset, touch done button, reward). We included only single-stroke trials. We first generated a “median trial”. For each segment (i.e., time window between a pair of successive anchor events), we found its median duration, and then concatenated these median segments to construct a median trial. We then aligned each trial to this median trial at the anchor events, warping time linearly within each segment. To avoid sharp discontinuities at anchor points, we smoothed the final firing rates at the times of the anchor points (2.5 ms Gaussian kernel). This warping did not change the firing rate values, just their timing.

Sorting of units (rows) in the resulting firing rate plot (Fig. 3c) was performed in cross-validated manner. Sort indices were determined using one subset of trials (n = 50) and then applied to sort the remaining subset of trials that are plotted (n = 235).

### Neural data analyses

#### Dimensionality reduction of population activity

##### Principal components analysis

We performed dimensionality reduction on the neural population activity, in general because high-dimensional noise can reduce the interpretability of the Euclidean distance^132^, and in one case in order to identify a potential linear projection of population activity (i.e., a subspace) that preferentially encodes primitives, a standard approach^37,133^. We represent a single area’s data from a single session and within a specific within-trial time window as a matrix X, of size N x KT, where N, K, and T are the number of units, trials, and time bins, constructed by concatenating time bins from all trials along the second dimension. Data were first binned in time (0.15 s window, 0.02 s slide) before constructing this data matrix. We used principal components analysis (PCA), but instead of applying PCA on single-trial data X, we applied PCA on trial-averaged data X_*C*_, in order to minimize the influence of trial-by-trial variation (noise). X_*C*_ holds the mean activity for each trial condition, of size N_C_ x KT, where N_C_ is the number of unique conditions, and where the specific conditions depended on the experiment (see below). We performed PCA on X_*C*_, and retained the top eight principal components (PCs). The specific trial-averaged conditions used for identifying PCs were the following. For analysis of motor invariance (**Fig. 4**), the conditions were each unique primitive (averaging over location/size), which resulted in identifying PCs that preferentially encoded primitives if they exist in the dataset. For analysis of categorical structure (**Fig. 5**), PCA was performed separately for each morph set, and the conditions were the unique images (i.e., the two endpoint shapes plus the morphed shapes in between). For the analysis of primitive representational reuse in characters (**Fig. 6**), the conditions were each combination of primitive and task type (i.e., resulting in *num_primitives* x *2* conditions).

We performed PCA in a cross-validated manner, to ensure that it was not overfitting to noise. We partitioned trials into two subsets (in a stratified manner), one “training” set that was used only for identifying the PCs, and a “test” set that was projected onto these PCs and then used for all subsequent analyses. We performed 8 randomized train-test splits (including all downstream analyses) and averaged their results.

##### Representing time-varying population activity as a vector

For some analyses, we captured single-trial time-varying population activity during the planning epoch (0.05 to 0.6 sec after image onset) as a vector, which could then be visualized after dimensionality reduction (**Extended Data Fig. 7**) or used in decoding analyses (**Extended Data Fig. 12**). Starting from a region’s population data, represented as a K x N x T matrix, where K, N, and T are the numbers of trials, units, and time bins (with time binned using a 0.2 s window, 0.1 s slide), we concatenated the N channels’ length T vectors end-to-end to construct the data matrix *D*, of size K x NT, where each trial is represented by a vector of length N x T.

##### Nonlinear dimensionality reduction

To visualize population activity in two dimensions (**Extended Data Fig. 7**), we used the nonlinear dimensionality reduction method Uniform Manifold Approximation and Projection (UMAP), performed on the *D*, using values of 40 and 0.1 for the parameters *n_neighbors* and *min_dist*.

#### Computing "neural distance”

To quantify the similarity of population activity between two sets of trials, such as trials for conditions A and B (where A and B are specific values of task-relevant variables), we devised a “neural distance” metric. This metric has the useful property of being unbiased (so that the expected value of the neural distance between two sets of trials sampled from the same distribution is zero). Inspired by the “normalized distance” in Liu et al^134^, it is the average pairwise Euclidean distance across conditions A and B, minus the within-condition distances. This subtraction step ensures that the distance is unbiased (unlike the mean Euclidean distance, which is biased upwards^35^). In addition, the resulting distance is normalized by dividing by an upper-bound distance to normalize it between 0 and 1. Neural distance is defined as:

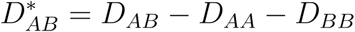

where the normalized Euclidean distance between sets of trial indices in conditions *A* and *B* is:

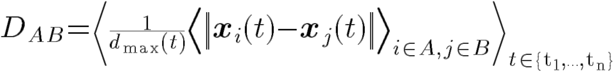

Here, *x_i_*(*t*) is the population activity vector at time *t* (in a window between times t_1_ and t_n_), and *d*_max_ is a normalization factor that is an upper-bound (98th percentile) of the distances between all pairs of different trials combined across all conditions.

#### Computing a variable’s encoding strength

To compute how strongly a given variable is encoded in population activity (e.g., “primitive encoding” in **Fig. 4j**), we computed the mean effect of that variable on population activity, in terms of neural distance, while controlling for the other relevant variables. Consider an experiment varying two variables, primitive and location, such that a condition is represented by the tuple (*p*, *l*), where *p* and *l* index the primitives and locations. Primitive encoding is the average neural distance across all pairs of conditions that have different primitives but same locations:

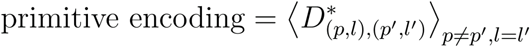

Location encoding is defined analogously:

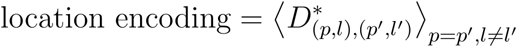

This approach generalizes to any pair of variables, such as primitive and task type in **Fig. 6**.

#### Statistically comparing brain regions in strength of variable encoding

In analyses that compare the encoding strengths of a particular pair of variables (e.g. primitive vs. location in **Fig. 4j**), we performed the following statistical tests to compare each brain region with every other brain region, in terms of how strongly they encode these two variables. We used the following procedure. (1) For each variable and pair of brain regions, we performed a statistical test comparing how strongly these two regions encoded that variable. This involved first extracting a dataset of neural distances between each pair of trial conditions for each of the two brain regions. For example, if the variable was “primitive”, then each of the two brain regions would contribute a dataset consisting of neural distance scores between all pairs of trial conditions that have different primitives but the same location. The datasets for these two regions would be combined into a single dataset, to which we fit a linear least-squares regression model to test for an effect of brain region on neural distance *y*, controlling for trial-condition pair:

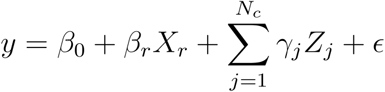

where *X_r_* is 0 or 1 depending on brain region, and *Z_j_* is an indicator variable for trial-pair condition, with *γ_j_* as their coefficients, and *ϵ* is a noise term. Finally, we extracted the p-value for *β_r_* (two-sided t-test), which represents the significance of the difference between this region pair in how strongly they encode the variable being tested. (2) This procedure was performed once for each combination of variable (2) and brain region pair (28), resulting in 56 p-values. We corrected the p-values for multiple comparisons using the Bonferroni method (28 brain region pairs x 2 variables = 56 comparisons). (3) Using these 56 values, we then summarized each region with two numbers representing the number of regions for which this region more strongly encodes these two variables. For example, for primitive x location experiments, each region was scored with a tuple (*N_prim_*, *N_loc_*), where *N_prim_* is the number of other regions that this region beats in the pairwise statistical tests of primitive encoding (and analogous for *N_loc_*, except that it scores location encoding). In summary plots (**Figs. 4j, 6f**), the results of this entire procedure were represented by mapping each region’s resulting tuple to a color based on a 2D color map.

#### Specific analyses

##### Analysis of motor invariance in neural activity

Dimensionality reduction was performed as described above, using a time window of 0.05 s to 0.6 s after image onset (to avoid including data after the go cue, which, for a subset of these motor-invariance experiments, occurred 0.6 to 1.0 s after image onset) for fitting the PCs and for analyses that involve time-averaging (**Fig. 4i-k**).

To test cross-condition decoder generalization (**Fig. 4k**), we used a linear support vector machine classifier (SVC), using a one-vs-the-rest scheme for multi-class classification (LinearSVC, scikit-learn, regularization parameter C set to 0.1). We report test accuracy linearly rescaled so that chance level (inverse of the number of classes) and 1 were mapped to 0 and 1. Because decoders were trained and tested on different conditions (with non-overlapping sets of trials), there was no concern of overfitting. Decoding was performed separately for, and averaged across, time bins (0.05 to 0.6 s relative to image onset).

##### Analysis of categorical structure in neural activity

Dimensionality reduction was performed as described above, using a time window from 0.05 s to 0.9 s after image onset for fitting the PCs. For analyses involving time-averaging (**Fig. 5d-g**), we used a window late in the planning epoch (0.6 to 1.0 s), when separation for A1 and A2 trials was the greatest (**Fig. 5h**). Primitive bias index was computed as above, using the Euclidean distance.

To assess whether primitives were associated with distinct activity patterns, we asked whether each primitive’s single-trial activity was separable from every other primitive’s activity, using a decoding approach (**Extended Data Fig. 12**). We used given region’s population activity during the planning epoch (0.05 to 0.6 sec after image onset) represented as a dataset *D*(size K x NT, where each trial is represented by a vector of length N x T; see section “Representing time-varying population activity as a vector”), with its dimensionality further reduced to K x 50 by performing PCA and keeping the top 50 principal components. For each pair of primitives decoding was performed using a linear support vector machine classifier (SVC) (LinearSVC, scikit-learn, regularization parameter C set to 0.1). Decoding was performed in a cross-condition manner to assess generalizable encoding of primitives; on each iteration we trained a decoder using data for drawings at one screen location (or size, for sessions with size variation), and tested using held-out data from all other locations (or sizes), and then averaging over analyses for different locations (or sizes). Because decoding performance can be biased upwards for regions with more units, to fairly compare the regions that had the strongest primitive encoding (preSMA, SMA, PMv, PMd, and vlPFC), we first matched their numbers of units by randomly subsampling (without replacement) to match the area with the fewest units and averaged the results from repeating this 10 times. By performing this overall procedure for every pair of primitives and separately for every brain region, each time returning a single decoding accuracy score, we populated the matrices in **Extended Data Fig. 12**. We combined results across five sessions for each subject (S1: 2 varying in location, 3 varying in size; S2: 3 location, 2 size).

##### Analysis for recombination of primitive representations in the Character task

For each session, we presented both Single-shape and Character tasks using a block-interleaved design (except one session using a trial-interleaved design). We analyzed primitives that were performed in both Single-shape (instructed by the shape image) and Character trials (the subject’s choice). For one experiment, the subject performed Multi-shape but not Single-shape tasks—we therefore used the first stroke from Multi-shape trials instead of Single-shape trials. For Character trials, we used only strokes that were scored as high-quality primitive matches. Dimensionality reduction was performed as above, using a time window of -0.8 to 0.3 s relative to stroke onset for fitting the PCs. For analyses involving time-averaging, we used a window -0.5 to -0.05 s relative to stroke onset. Neural distance, primitive encoding, and task-type encoding were computed as above, using primitive and task type (instead of location or size) as the two relevant variables.

For analyses comparing strokes from the Single-shape task to the non-first stroke of the Character and Multi-shape tasks (i.e., the second stroke and onwards) (**Extended Data Fig. 14**), all parameters were the same as in the main analysis comparing Single-shape strokes to the first stroke of Character drawings, except two differences. First, we used a shorter analysis time window of -0.35 to -0.05 s relative to stroke onset (instead of -0.5 to -0.05 s). This was to account for the duration of the gap between offset of one stroke and onset of the next (mean of ∼0.3 sec) in Character drawings. Second, we preprocessed the neural data to account for the presence of a strong effect of the reaching movement to initiate the trial (i.e., reaching from the “Hold button” towards the image; see **Fig. 1c**), which was present for Single-shape trials, but not for the non-first strokes of Character and Multi-shape trials. This effect is clear in population trajectories (**Extended Data Fig. 14a**). To accurately assess whether primitive-encoding activity was reused across task types, it was important to correct for this effect of initial reach, because it was present for Single-shape trials, but not for the non-first strokes of Character and Multi-shape tasks. We performed this correction in the simple manner, by subtracting, for all strokes, the across-stroke mean effect of the initial reach. This correction was performed separately for each time bin as follows. For each unit, we first estimated the mean effect of initial reach by using linear least-squares regression. Activity for each stroke was modeled as:

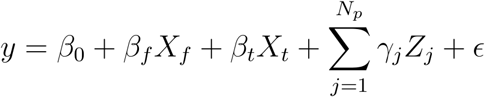

Here, is *y* is firing rate, *X_f_* is 0 or 1 depending on whether the stroke is the first stroke (always 1 for Single-shape data), *X_t_* is 0 or 1 depending on task type (Single-shape or Character), *Z_j_* is an indicator variable for the primitive class, and *ϵ* is a noise term. To correct for the effect of initial reach, we subtracted *β_f_* (the mean effect of initial reach) from the neural activity for all first strokes. We did not perform this correction for comparisons of Single-shape strokes to the first stroke in Character drawings (**Fig. 6**), because, in those cases, all strokes included the initial reaching movement.

##### Analysis of primitive encoding aligned to visual fixations

We assessed the extent to which PMv activity, aligned to visual fixation events during the Multi-shape task, preferentially encodes either the primitive associated with the visually-fixated shape or the first primitive the subject is planning to draw (**Extended Data Fig. 15**). We first converted time-varying eye tracking data (x- and y-position time series) to a sequence of fixation and saccade events. This was done using the Cluster Fix algorithm^135^, which, briefly, uses k-means clustering on distance, velocity, acceleration, and angular velocity, and then assigns clusters to fixation and saccade events. Each fixation event was assigned a “visually-fixated” primitive based on the closest shape. If all shapes were further than 70 pixels (2.08 cm), then this fixation event was considered to be looking away from all shapes and was thus excluded from further analysis. Each fixation event was assigned a “planned primitive” based on which primitive the subject would go on to draw first on that trial. Note that choices were made freely, without experimenter-imposed rules.

We assessed the extent to which fixation-aligned neural activity encoded each primitive using a decoding approach, where the strength of representation of a given primitive was defined as the probability score returned by a decoder trained to classify that primitive vs. all other primitives. Decoders were trained using Single-shape trials from the same session. Data from the planning epoch of Single-shape trials were “cut” into multiple short-duration “snippets”, each of which was a datapoint used for training the decoder. Specifically, neural data was represented as a matrix X, of size N x KT, where N, K, and T are the number of units, trials, and time bins, constructed by concatenating time bins from all trials along the second dimension (data were first binned in time using a 0.3 s window with 0.1 s slide) before constructing this data matrix. These KT N-dimensional training data points, each with an associated “primitive” label, were used to train a multilabel, logistic regression, one-vs-rest classifier (*scikit-learn*), which pools multiple independent classifiers, one per primitive. This resulted in a multilabel decoder that could be applied to neural data at any time point and return one probability score for each primitive, where the probability can be interpreted as the strength of that primitive’s representation.

##### Analysis of encoding of kinematic variables in neural activity

To test for encoding of generic kinematics in neural activity (**Extended Data Fig. 16**), we used a standard encoding model approach, based on a recent study of encoding of handwriting kinematics in human motor cortex^55^. We assessed the fraction of variance in neural activity that is explained by a linear mapping from moment-by-moment finger kinematics to neural activity. As in that human study, we used neural activity in the top 10 dimensions found after performing PCA on the neural activity, which helped to reduce the effect of noise present in higher dimensions. PCA was performed as we described above, except, instead of identifying the PCs using trial-averaged data, we did so using single-trial data, in order to retain activity potentially related to trial-by-trial variation in kinematics. Activity at each time point was modeled as:

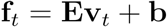

Here, f_*t*_ is the neural activity at time bin *t* in the top 10 neural PCs, E is a 10 x 2 matrix mapping kinematics to neural activity (i.e., preferred directions), v_*t*_ is the 2 x 1 finger velocity and b is an intercept term. The fraction of variance accounted for (FVAF) was computed as follows:

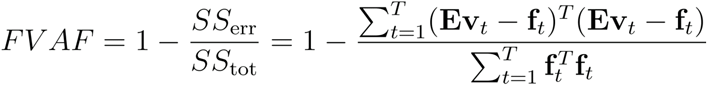

Here, *SS*_tot_ is the total variance, and *SS*_err_ is the sum of squared errors, and *T* is the total number of time steps across all trials.

To test how strongly neural activity encoded kinematics in a manner that generalized across primitives, the model was trained on one subset of primitives (all except one) and tested on one held-out primitive. This was performed once for each primitive, with the final FVAF computed as the mean FVAF across all train-test splits. We used sessions performing the Single-shape task with location and size variation (S1, n = 30 primitives across five sessions; S2, n = 42 primitives across three sessions, which excludes two sessions with fewer than six primitives). We included data throughout each stroke’s duration.

To allow for the possibility that kinematics better relates to neural activity at a non-zero lag, we performed this analysis multiple times with different time lags between neural and behavioral data, varying from -0.3 to 0.3 s (0.05 s increments). The final scalar summary averaged the results in a time window from -0.15 s to -0.05 s (with neural leading behavior), consistent with a peak time lag of around -0.1 s (neural leading) found in prior studies^136^ and in our analysis (**Extended Data Fig. 16**).

Note that the FVAF values we found in M1 are comparable to the average value of 0.3 in the prior study of motor cortex in human handwriting^55^. Our finding of lower values (ranging from around 0.0 to 0.2, **Extended Data Fig. 16b**) is consistent with our use of trial-level, instead of trial-averaged data—which introduces more variability—and our approach of testing generalization across primitives (if we use a more traditional method of holding out trials, and not primitives, we find FVAF values are higher, with values ranging from around 0.1 to 0.3).

## Acknowledgements

We thank Y. Liu, V. Goudar, X. Ma, T. Wu, D. Dolnik, S. Coolsaet, S. Sharma, A. Urquieta, V. Calligy, and other members of the Freiwald, Wang, and Tenenbaum labs for project feedback; F. Buck, D. Hildebrand, E. Hosseini, P. Jaffe, K. Kay, T. Nigam, Y. Prut, J. Rhee, P. Schade, T. Suzuki, and W. Zarco for manuscript feedback; V. Sherman and A. Gonzalez for technical assistance; and L. Ying for administrative assistance. This work was supported by the National Institutes of Health, through the National Eye Institute (R01EY021594 to W.A.F.), the National Institute Of Mental Health (F32MH125573 to L.Y.T.), and the National Institute Of Neurological Disorders And Stroke (K99NS131585 to L.Y.T.), as well as the Simons Foundation’s Collaboration on the Global Brain (876120SPI and AN-NC-GB-Pilot Extension-00002596-01 to X.-J.W., J.B.T., and W.A.F., and NC-GB-CULM-00003138 to X.-J.W.), the Center for Brains, Minds & Machines of the National Science Foundation (STC award CCF-1231216 to W.A.F. and J.B.T.), the Office of Naval Research (N00014-20-1-2292 to W.A.F., N00014-23-1-2040 to X.-J.W., and MURI N00014-21-1-2801 to J.B.T.), and the Air Force Office of Scientific Research (FA9550-22-1-0387 to J.B.T.).

## Author contributions

Study conception and design: L.Y.T., X.-J.W., J.B.T., and W.A.F. Surgery: L.Y.T., A.G.R., M.A.G.E., M.H.S., and W.A.F, with input from K.U.G. Experiments: L.Y.T., with input from K.U.G. Data analysis: L.Y.T., with input from K.U.G. and D.J.H. Manuscript initial draft: L.Y.T. Manuscript revision: L.Y.T., X.-J.W., J.B.T., and W.A.F., with input from the other authors. Funding acquisition: L.Y.T., X.-J.W., J.B.T., and W.A.F. Project supervision: X.-J.W., J.B.T., and W.A.F.

## Data and code availability

The data and code used in this study will be made available on Zenodo upon publication.

---

**Extended Data Fig. 1.**
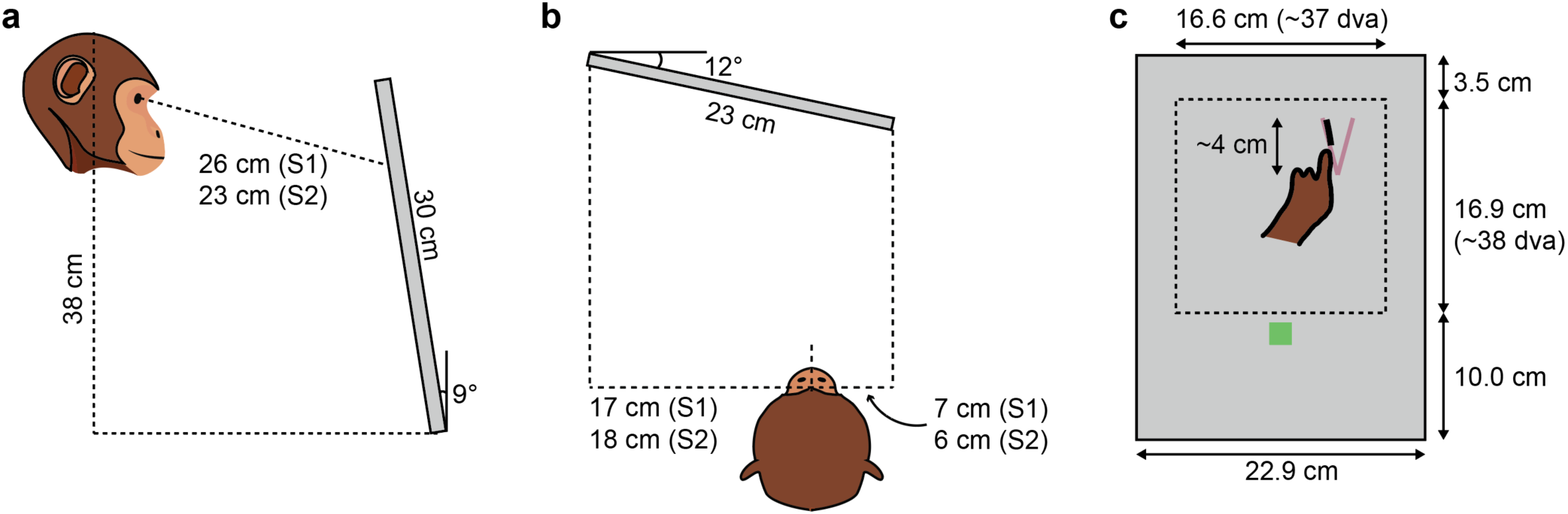
Behavioral task setup. (a) Schematic of subject relative to screen, profile view. Subjects 1 and 2 were positioned at different distances to accommodate their individual anatomies and postures. The screen was slanted slightly to optimize the ability to see and reach to the same part of the screen (the workspace at the top of the screen). (b) Schematic of subject position relative to screen, top view. Subjects were positioned to the right, to accommodate reaching to the screen with the hand used for drawing (left). (c) Schematic of screen during trial, with component locations and sizes to scale. The finger is tracing over the figure (purple-gray “V”), leaving a black trail of “ink” behind. The average size of shapes was 4.0 cm (maximum of width and height). The thickness and color of the figure and “ink” are to scale. The “done” button is visible (green square), and the subject can press it at any time to report completion. The dashed line indicates the workspace (not visible to the subject).

**Extended Data Fig. 2.**
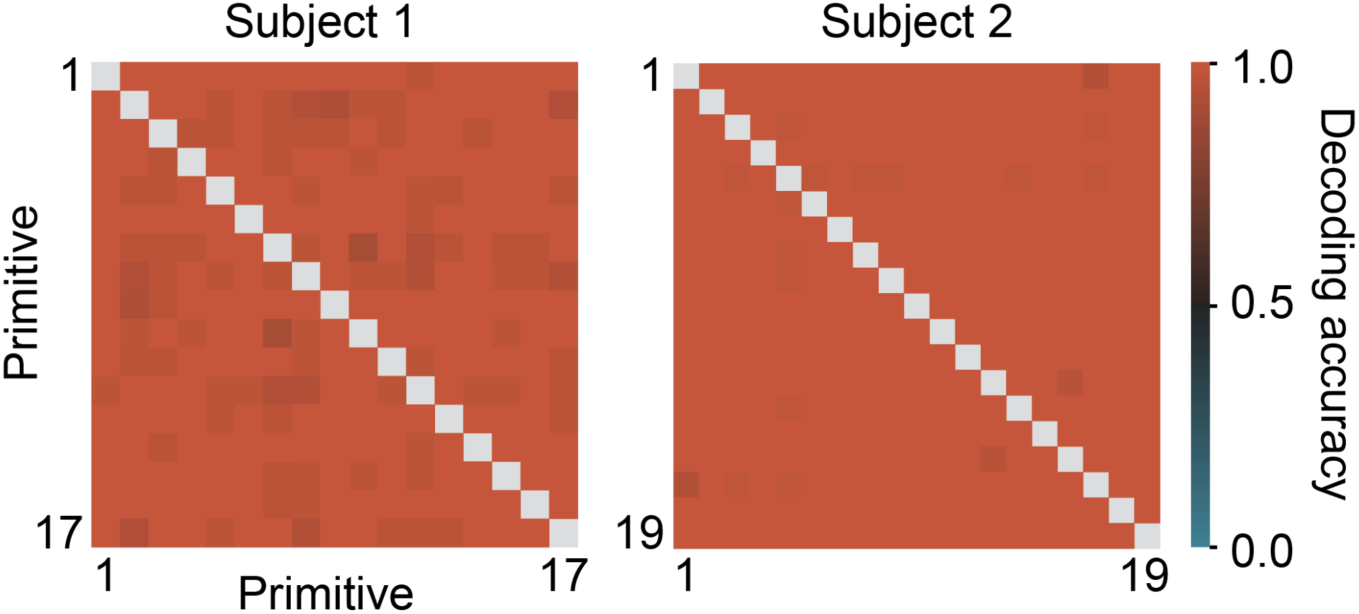
Primitives are distinct from each other in their single-trial kinematics. Pairwise decoding accuracy between all pairs of primitives, using stroke single-trial kinematics. Strokes were represented as 8D vectors derived from their x- and y-position time series (see Methods). Cross-validated decoding was performed separately for each pair of primitives, and the results for all pairs are combined in this summary matrix. Chance decoding is 0.5, because each analysis had two primitive classes. Decoding accuracy was high for all primitive pairs (minimum accuracy was 0.90 for S1 and 0.94 for S2).

**Extended Data Fig. 3.**
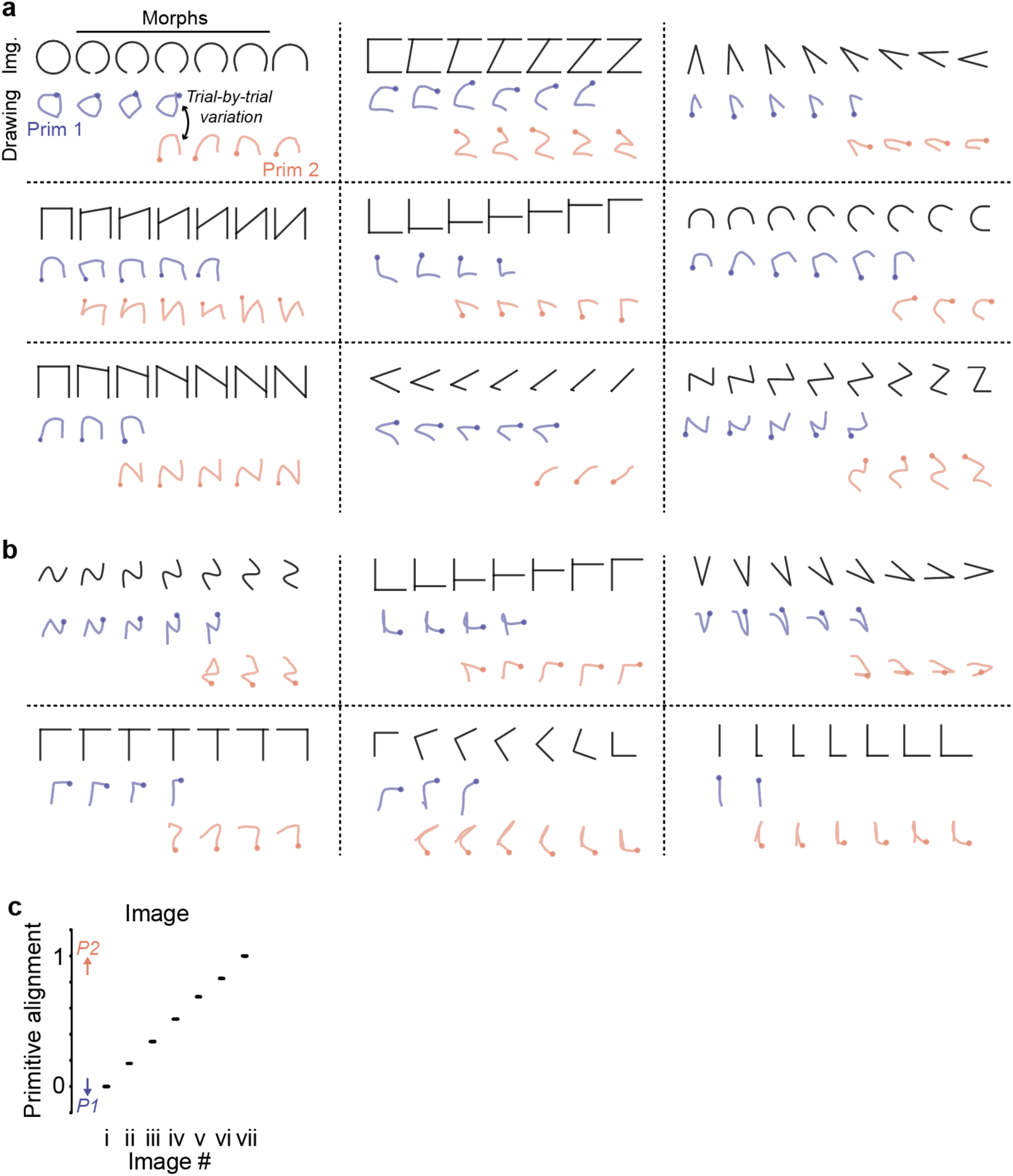
Categorical structure in behavior: more examples and quantification of primitive alignment for image data. (a) Example single-trial drawings across nine different morph sets, for subject 2. Drawings are colored by whether they reflect use of primitive 1 (blue) or primitive 2 (orange). Images morph between two well-practiced shapes. The examples here are depicted in a similar manner to the example in Fig. 2. (b) Same as panel a, but for subject 1. (c) Primitive alignment vs. trial condition for the example experiment in panel Fig. 2b, performed on image data (using the image distance), and not on behavioral data as in Fig. 2i.

**Extended Data Fig. 4.**
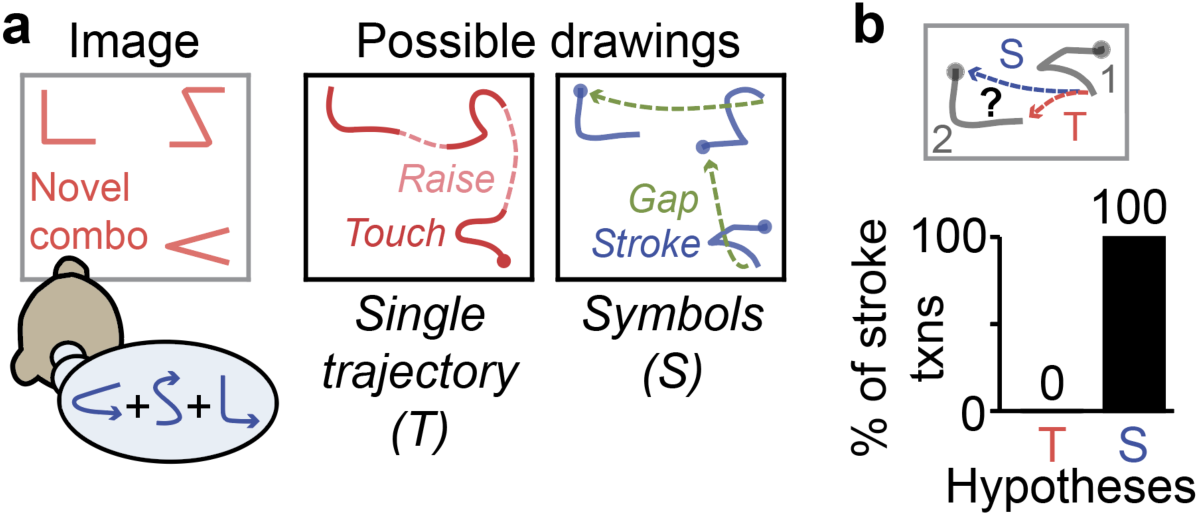
Recombination of stroke primitives into sequences, in the Multi-shape task. (a) Experiment testing for recombination of primitives into sequences, using the Multi-shape task. Given images composed of multiple disconnected shapes, two possible drawing responses are shown, consistent with either the “Single trajectory” (T) or “Symbols” (S) hypotheses (see main text). (b) Fraction of stroke-to-stroke transitions in which the second stroke is drawn in a manner consistent with the Single trajectory (T) or Symbols (S) strategies, restricted to transitions where the behavioral predictions of these two strategies differed [bottom bar plot, n = 391 transitions (S1), 529 (S2)]. Top, schematic of an example transition between two strokes labeled 1 and 2. Stroke 2 can be drawn either starting from the top-left, consistent with primitive reuse (“Symbols”), or from the bottom right, which would reflect the subject taking the shorter of the two gap distances (“Single trajectory”).

**Extended Data Fig. 5.**
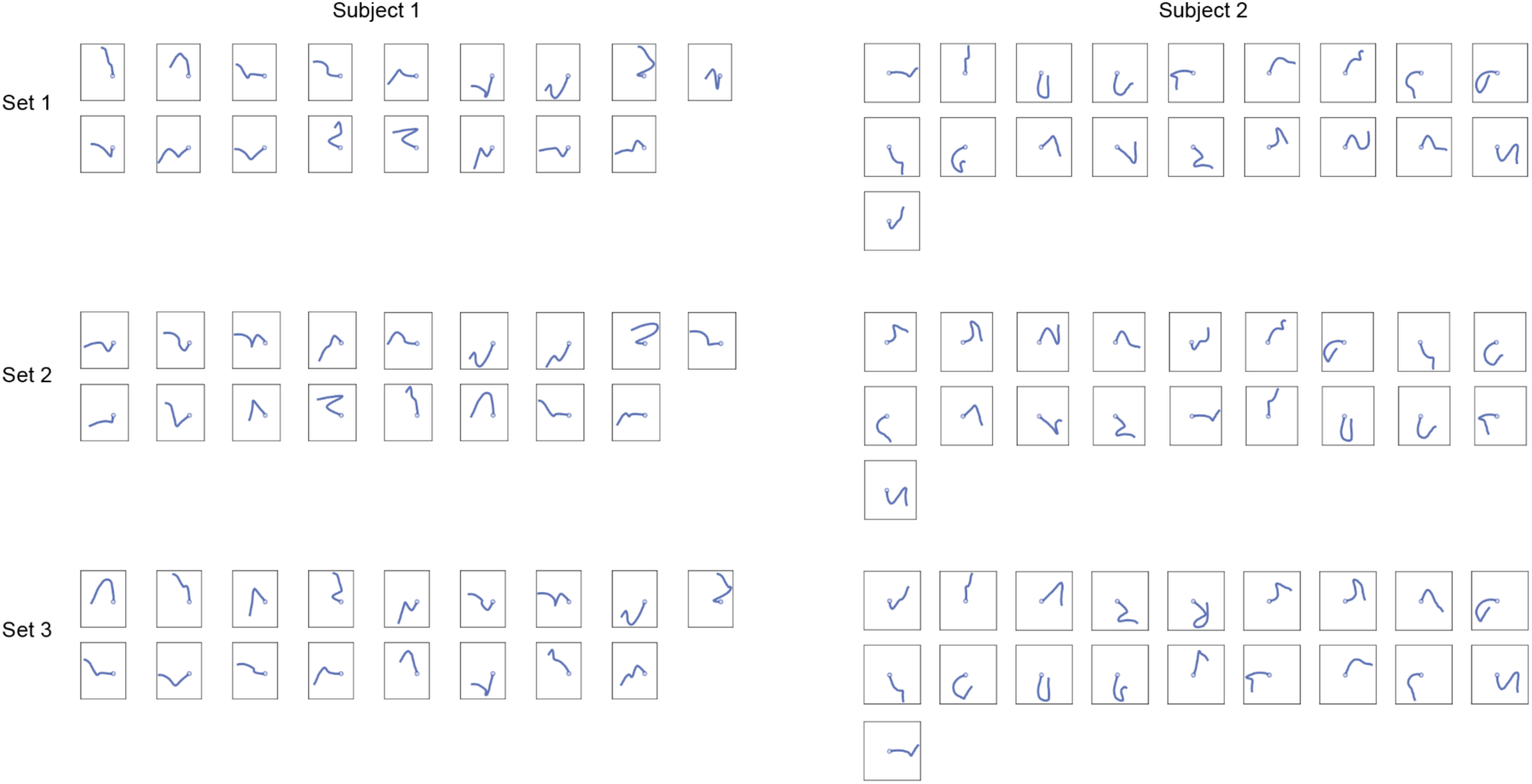
Simulated primitives created by “remixing” each subject’s actual primitives. Three sets of shuffled primitives for each subject are shown. Each shuffled primitive was constructed by connecting the first half of one primitive to the second half of another primitive drawn by the same subject. Each set was optimized so that shuffled primitives within a set were sufficiently different from each other and were also sufficiently different from all actual primitives. See Methods for details. These primitives were used in Fig. 2p.

**Extended Data Fig. 6.**
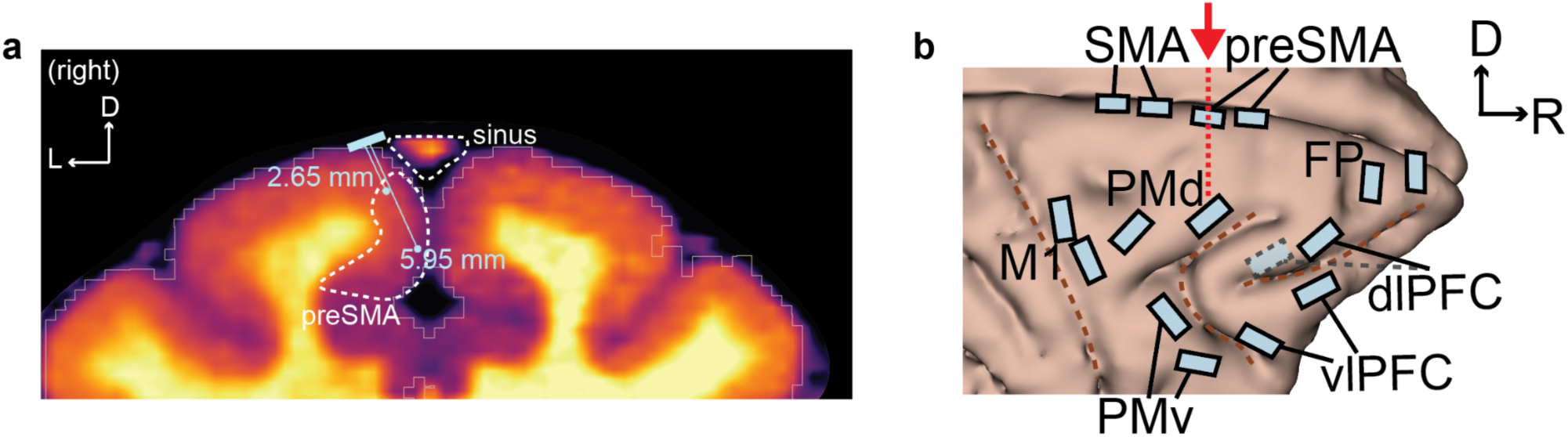
Angled implantation of SMA and preSMA arrays to avoid the sinus and target the medial wall. (a) Coronal MRI section showing the surgical plan for location and angle of a preSMA array relative to the superior sagittal sinus (“sinus”). SMA and preSMA arrays were implanted ∼2 mm lateral to the midline, and angled medially, to target the medial wall (preSMA), while avoiding the superior sagittal sinus. The array is depicted as a rectangular blue surface and two lines representing the shortest (2.65 mm) and longest (5.95 mm) electrodes. This location and angle is our best estimate based on the stereotactic coordinates and intra-operative photographs. The other two SMA and one preSMA arrays were located and angled similarly. D, dorsal. L, lateral. (b) Rostral-caudal location of the coronal section in panel a (red arrow and dashed line), overlaid on a brain surface model. D, dorsal. R, rostral.

**Extended Data Fig. 7.**
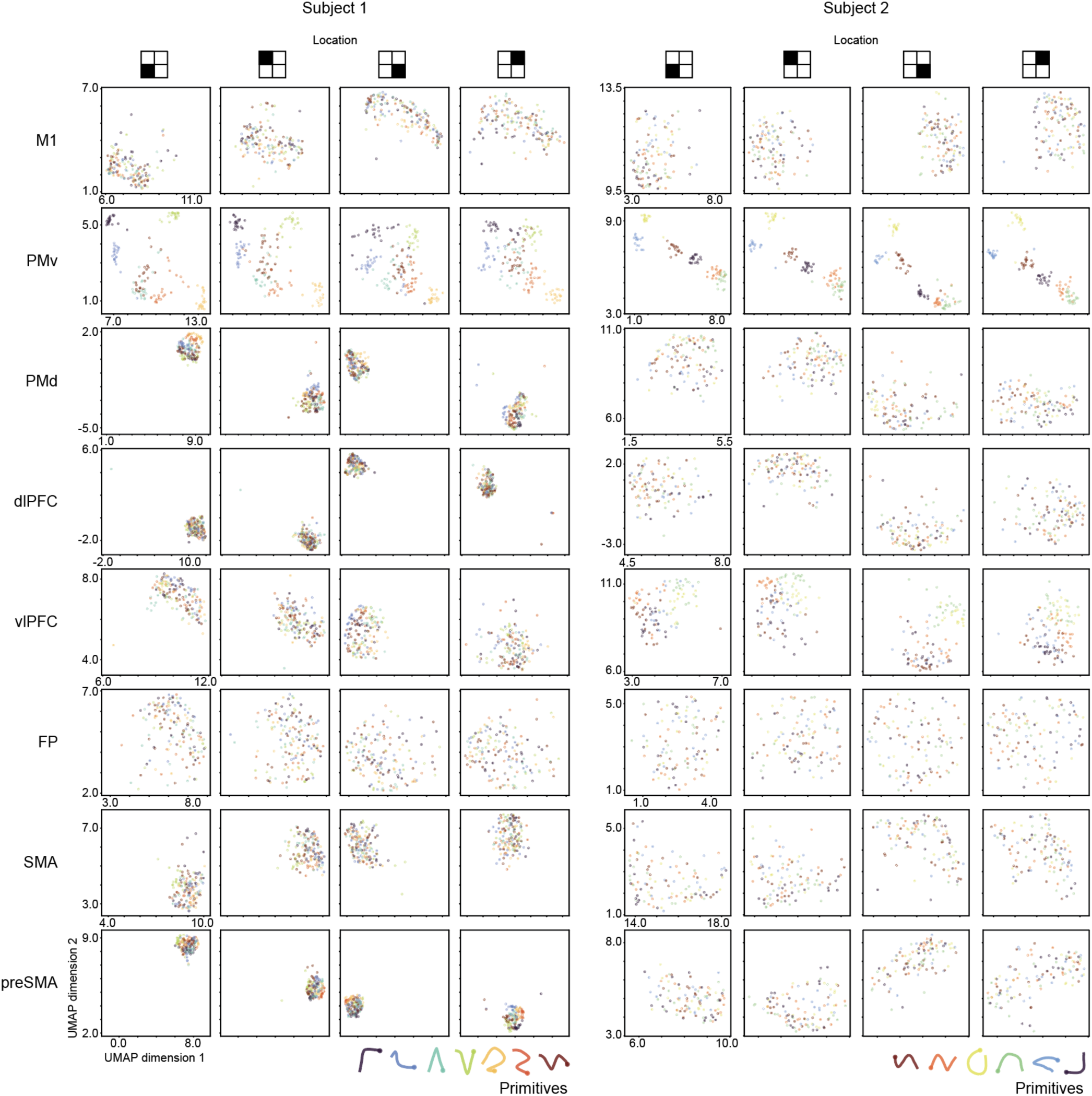
Separation of single-trial activity by primitive, invariant to location. Two-dimensional embedding of single-trial population activity using Uniform Manifold Approximation and Projection (UMAP). Each point represents a single trial, colored by primitive and split by brain region (rows) and screen location (columns). UMAP was performed on a vector representation of single-trial activity, constructed by concatenating all units’ time-varying activities into a single vector (see Methods). Activity was taken from the planning epoch (0.05 to 0.6 sec after image onset), from a single session of the Single-shape task, in which primitives varied in location. It is visually apparent that PMv activity clusters by primitive and that this clustering is largely unaffected by location. In contrast, activity in other regions is either unstructured (FP), varies strongly by location with minimal effect of primitive (M1, PMd, dlPFC, SMA, preSMA), or varies somewhat by primitive but in a manner that depends strongly on location (vlPFC).

**Extended Data Fig. 8.**
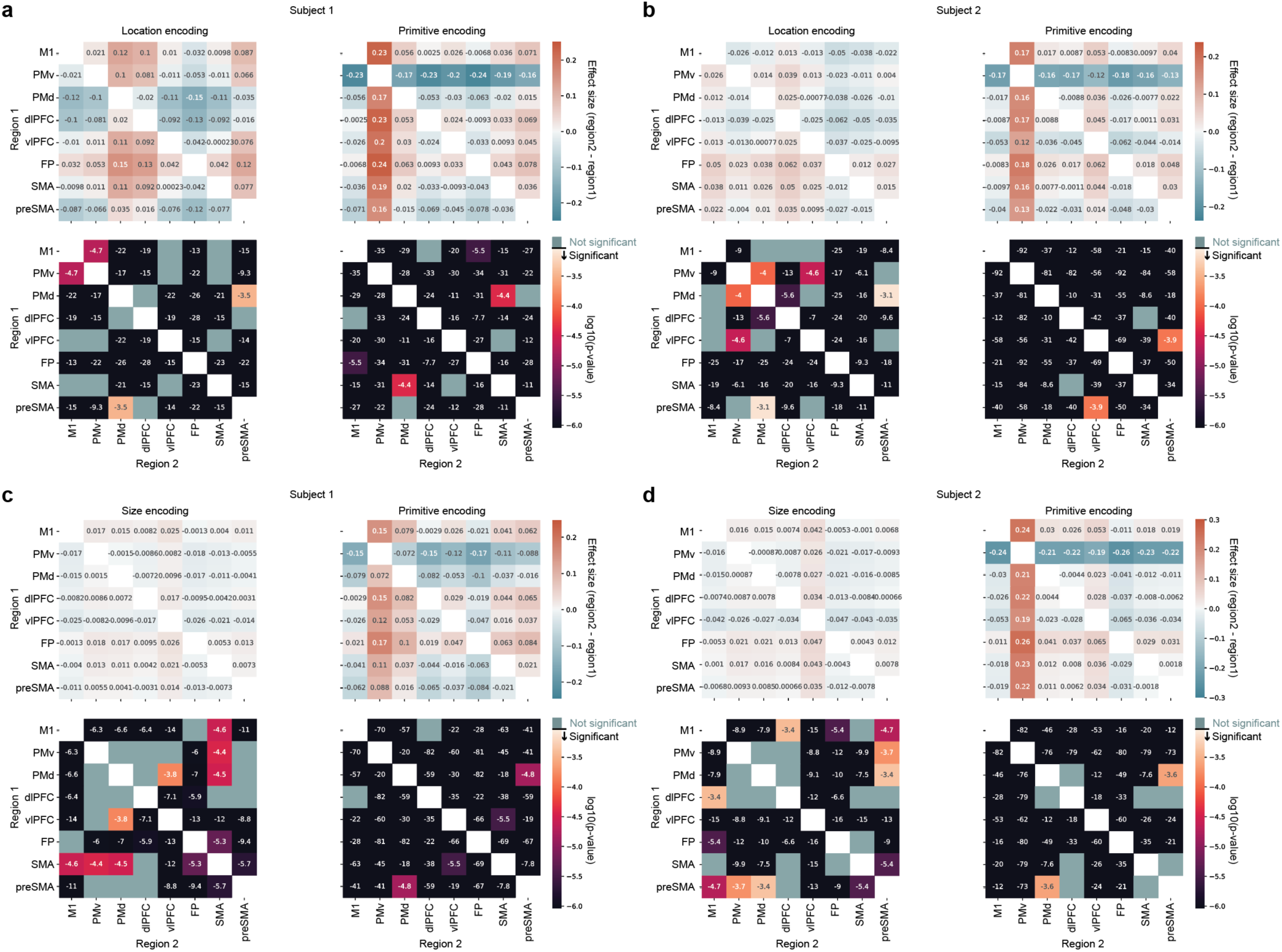
Detailed statistics for the analysis of location invariance in Fig. 4 and size invariance in Extended Data Fig. 10. (a) Effect size (top) and p-value (bottom) for comparisons between each pair of brain regions, for location encoding (left) and primitive encoding (right), for subject 1. Corresponds to Fig. 4j. “Significant” for the p-value heatmap means that p < 0.05 after Bonferroni correction for 56 comparisons (28 brain region pairs and 2 variables, location and primitive). (b) Same as panel a, but for subject 2. Corresponds to Fig. 4j (c) Same as panel a, but for size, instead of location, encoding. Corresponds to Extended Data Fig. 10b. (d) Same as panel c, but for subject 2. Corresponds to Extended Data Fig. 10b.

**Extended Data Fig. 9.**
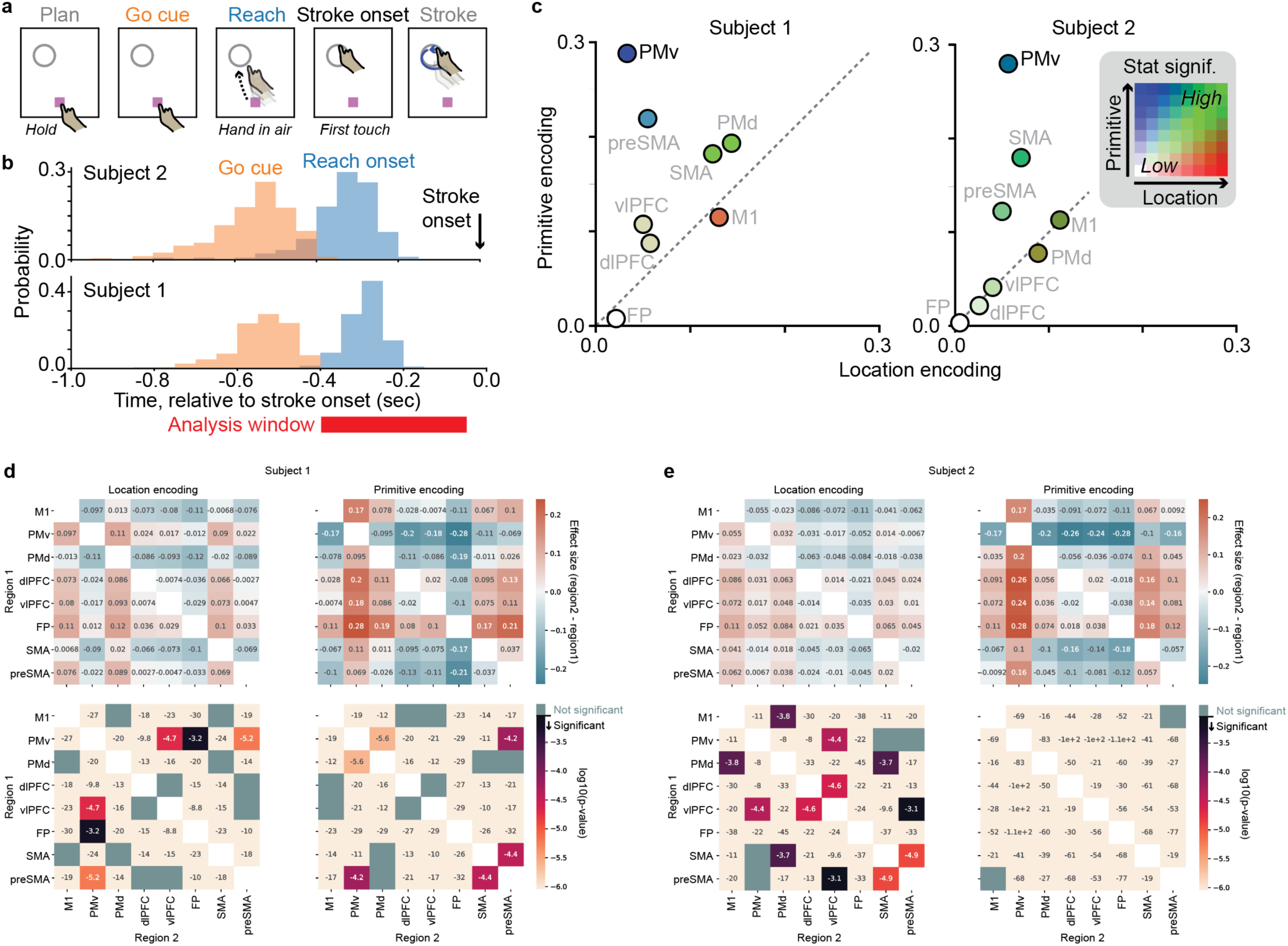
Invariant encoding of primitives in PMv during the initial reaching movement. (a) Schematic of trial events, highlighting the initial reaching movement from the location of the “hold” button to the location of the stroke onset. (b) Histogram of times of the go cue (orange) and reach onset (blue) relative to the time of stroke onset. To analyze activity during the initial reach, we used the time window -0.4 s to -0.05 s relative to stroke onset. (c) Summary of primitive encoding and location encoding across areas and sessions. Points show mean encoding scores. The color of each point denotes statistical significance (analogous to Fig. 4j). It indicates the number of other brain areas that this area “beats” in pairwise statistical tests of primitive encoding and location encoding (represented in the inset heatmap, “low” to “high” ranges from no (0) to all (7) other regions beaten). For statistics, each data point was a unique pair of primitive-location conditions (trial-averaged). For testing primitive encoding, these pairs were the same location, but different primitive [n = 93 (subject 1), 288 (subject 2)]. For location encoding, the pairs were the same primitive, but different location [n = 114 (S1), 132 (S2)]. Statistical tests were performed on condition pairs pooled across sessions (2 for S1, 3 for S2). (d) Statistics for primitive and location encoding in panel c. Effect size (top) and p-value (bottom) for comparisons between each pair of brain regions, for location encoding (left) and primitive encoding (right), for subject 1. “Significant” for the p-value heatmap means that p < 0.05 after Bonferroni correction for 56 comparisons [28 brain region pairs and 2 variables (location and primitive)]. (e) Same as panel d, but for subject 2.

**Extended Data Fig. 10.**
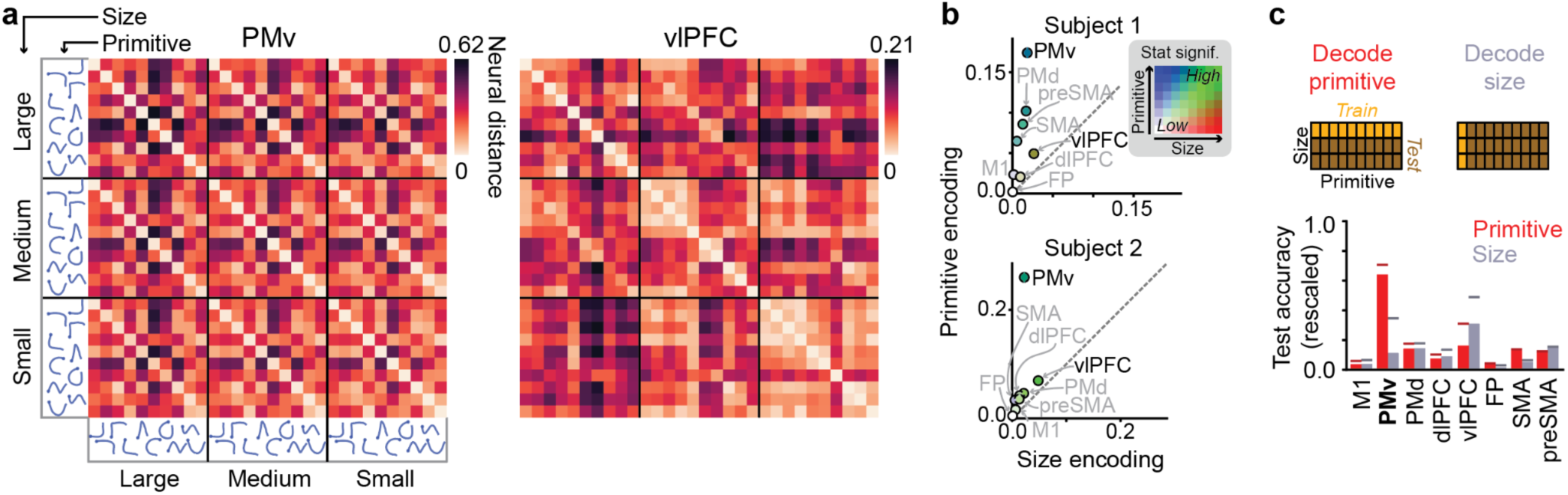
Size-invariant encoding of stroke primitives in PMv. (a) Matrix of pairwise neural distances between each unique combination of primitive and size, averaged over the planning epoch (0.05 to 0.6 s relative to image onset), for PMv and vlPFC. Shown are data from a single session for subject 2. n = 17 - 23 trials per primitive-size combination. (b) Summary of primitive encoding and size encoding across areas and sessions, with each point depicting the encoding scores for a given area. Each point’s color denotes statistical significance, in terms of the number of other brain areas that this area beats in pairwise statistical tests of primitive encoding and size encoding (represented in the inset heatmap; “low” to “high” ranges from no (0) to all (7) other regions “beaten”); see Fig. 4j and Methods for details. Each data point was a unique pair of primitive-size conditions (trial-averaged). For testing primitive encoding, these pairs were the same size, but different primitive [n = 313 (subject 1), 253 (subject 2)]. For size encoding, the pairs were the same primitive, but different size [n = 60 (S1), 56 (S2)]. Statistical tests were performed on condition pairs pooled across sessions (3 for S1, 2 for S2). For exact effect sizes and p-values, see Extended Data Fig. 8c, d. (c) Across-condition generalization of linear SVM decoders for primitive (red) and size (gray). See explanation in Fig. 4k.

**Extended Data Fig. 11.**
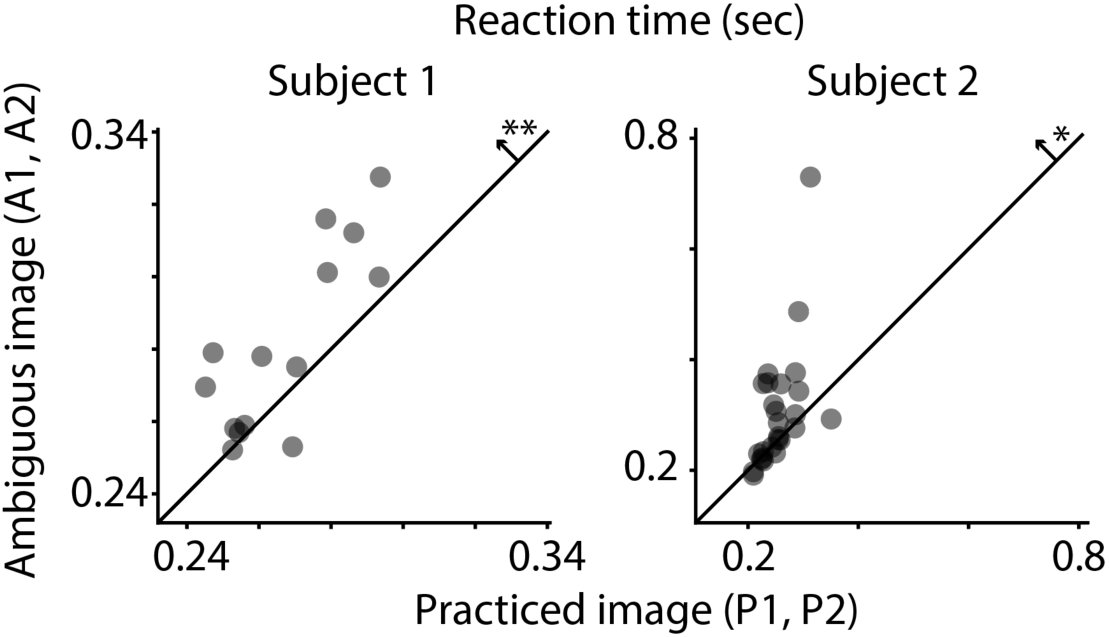
Slower reaction time for ambiguous images in tasks testing categorical structure. Average reaction time in seconds (between go cue and stroke onset), comparing ambiguous images to practiced images. Each data point represents a single primitive from one morph set, its y-value indicating reaction time when the it was drawn in response to an ambiguous image, and its x-value indicating reaction time when it was drawn in response to the practiced image. **, p = 0.0031, Wilcoxon signed-rank test (n = 14, W = 8); *, p = 0.024, Wilcoxon signed-rank test (n = 26, W = 87). Each data point is a primitive, and each morph set contributes two primitives (one at each morph endpoint).

**Extended Data Fig. 12.**
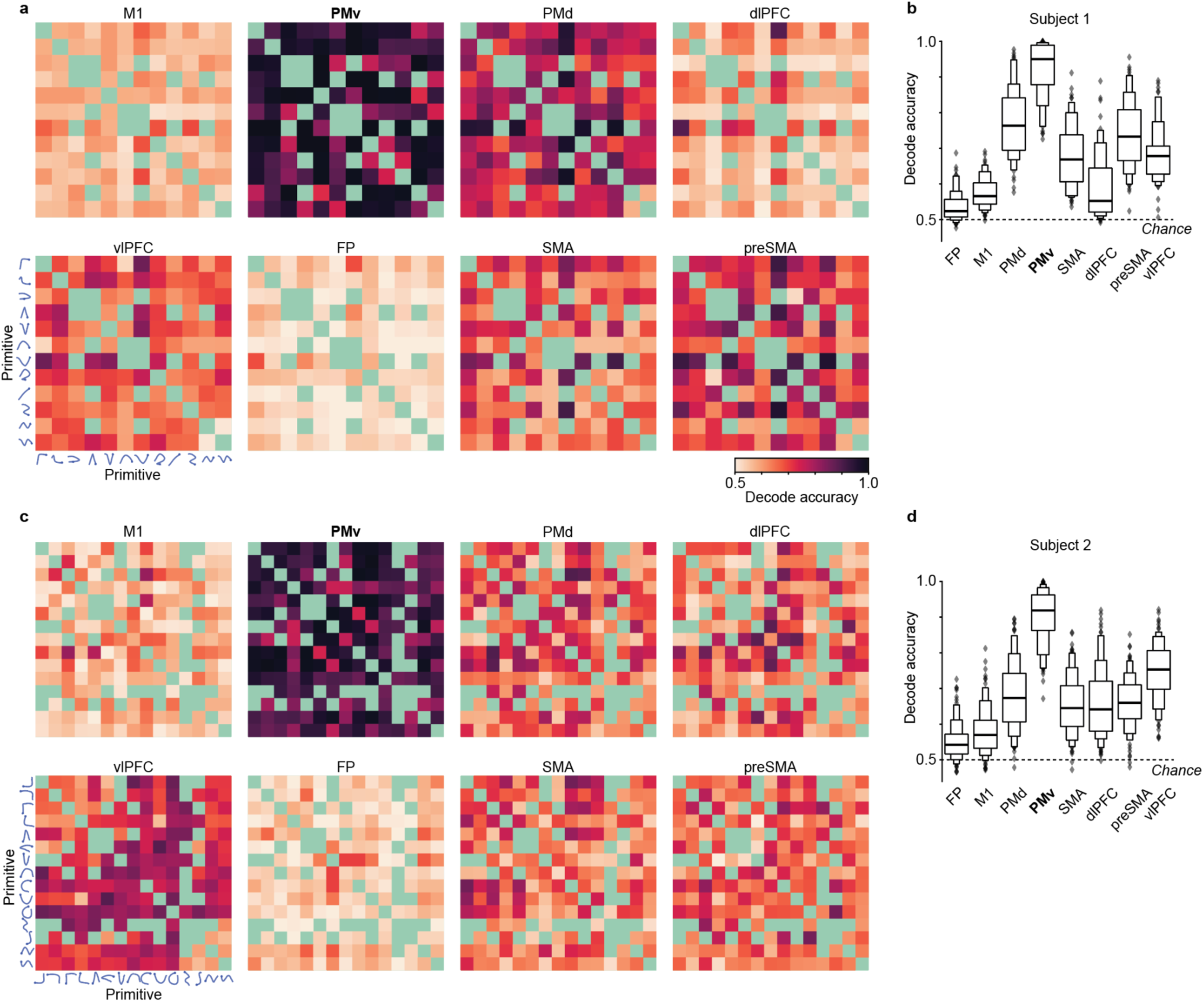
Primitives are distinct from each other in their planning-related activity in PMv. (a) Pairwise decoding between all pairs of primitives (those performed in the same session), for subject 1, using population activity from the planning epoch (0.05 to 0.6 sec after image onset) represented as a 50-dimensional vector (see Methods). Chance decoding is 0.5. The matrices aggregate across sessions (n = 5 Single-shape sessions for each subject; S1: 2 varying in location, 3 varying in size; S2: 3 location, 2 size). The gray-green cells are pairs which lack data because the primitives were not recorded in the same session. (b) Summary of decoding accuracy for all primitive pairs, for Subject 1, shown in a letter-value plot, a variation of a boxplot, in which each box depicts a successively decreasing quantile, from median, to quartile, to upper and lower eights, and so on. Each datapoint is one unique primitive pair. (c) Like panel a, but for subject 2. (d) Like panel b, but for subject 2.

**Extended Data Fig. 13.**
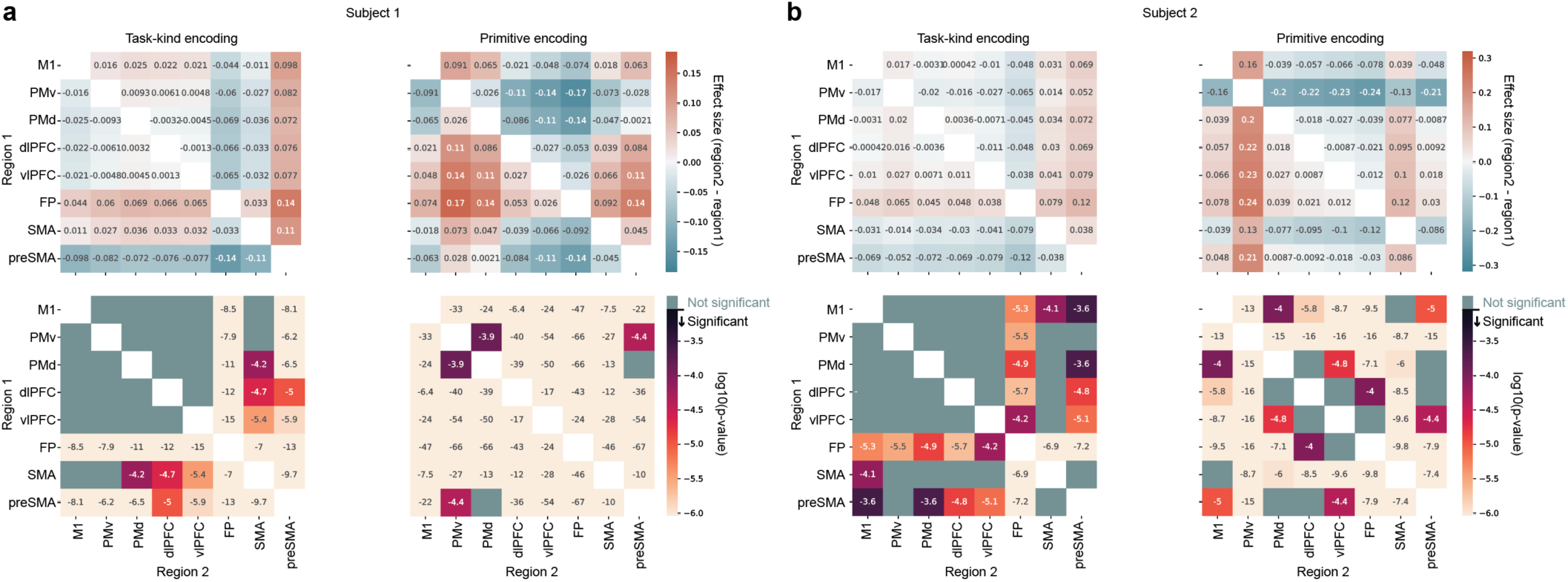
Detailed statistics for the analysis of recombination in Fig. 6. (a) Effect size (top) and p-value (bottom) for comparisons between each pair of brain regions, for task-type encoding (left) and primitive encoding (right), for subject 1. Corresponds to Fig. 6f. “Significant” for the p-value heatmap means that the value is less than 0.05 after Bonferroni correction for 56 comparisons [28 brain region pairs and 2 variables (task type and primitive)]. Note that because analyses involved bootstrapping to balance sample sizes across sessions (see Fig. 6f), these are statistics from a single bootstrapped sample that matches the mode. (b) Same as panel a, but for subject 2. Corresponds to Fig. 6f.

**Extended Data Fig. 14.**
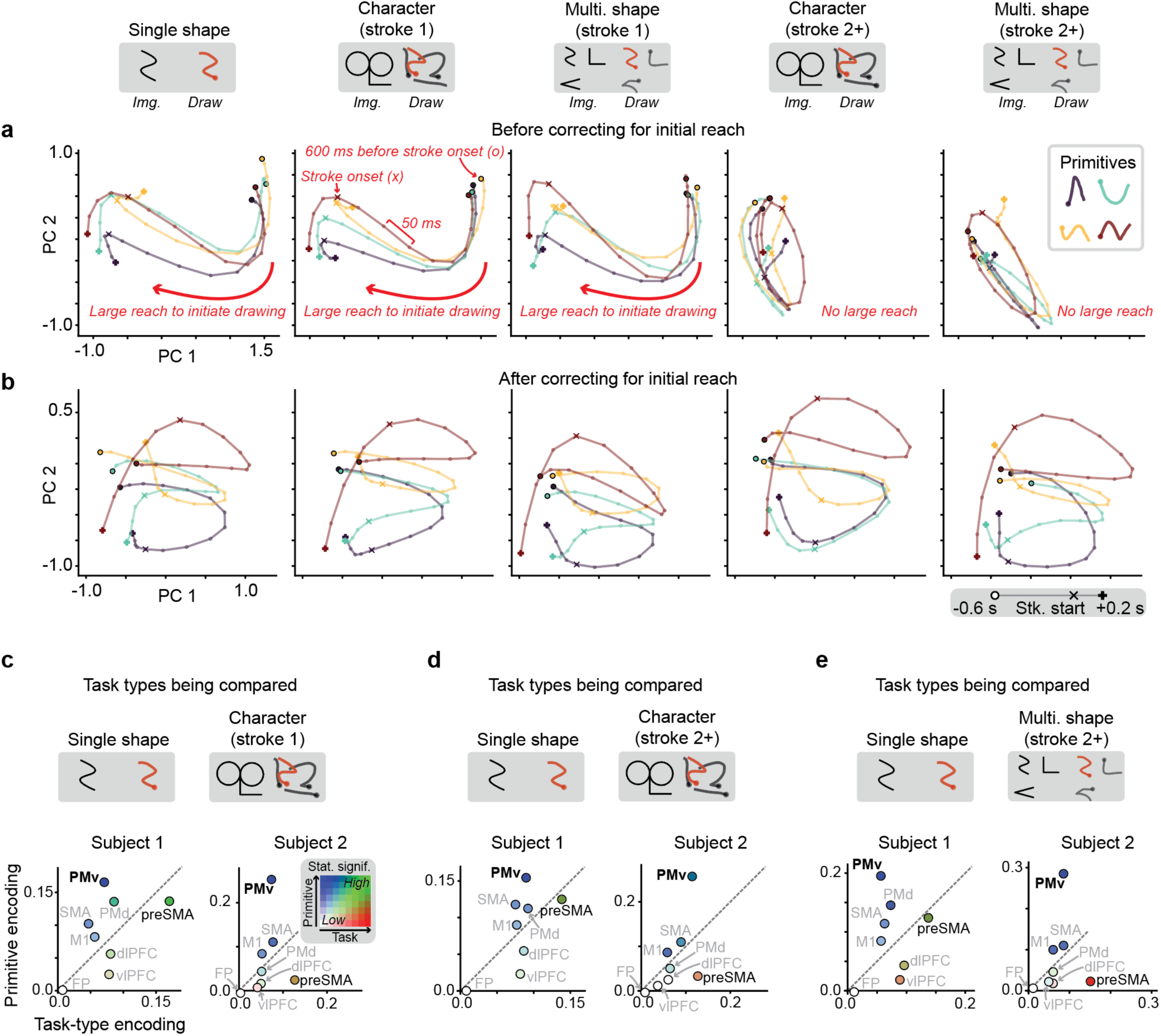
Recombination of primitives into sequences is reflected in PMv, including for the second stroke onwards (related to Fig. 6). (a) PMv population trajectories for each primitive (color), plotted in a subspace spanned by principal components 1 and 2, and aligned to stroke onset (“x” icon). Each subpanel shows a different subset of trials (i.e., strokes) from this session, differing by task type (Single-shape, Character, or Multi-shape) and whether the dataset includes only the first stroke or only the second stroke onwards (stroke 1 or stroke 2+). Note that the Single-shape task by design only has one stroke. A prominent activity pattern visible in these trajectory plots is the right-to-left sweep associated with the initial reaching movement to start the drawing. This activity is visible for the conditions that involve an initial reach—Single-shape, Character (stroke 1), and Multi-shape (stroke 1)—and not for conditions that lack this reach—Character (stroke 2+) and Multi-shape (stroke 2+). (b) The same as panel a, but after correcting for the effect of the initial reach. This correction was performed by subtracting, identically for all trials, the across-trial mean effect of the initial reach on neural activity, performed separately for each time bin (see Methods). After applying this simple linear correction, the primitive-associated trajectories are now similar across all conditions, consistent with the reach effect being an additive component (on top of the primitive-encoding component), associated with the large movement of the arm during the initial reach. (c) Summary of primitive encoding and task-type encoding across areas and sessions, comparing Single-shape to Character (stroke 1) trials, after correcting for the mean effect of initial reach on neural activity. Apart from this correction, this is the same data as in Fig. 6f. For panels c-e, the plotting conventions are identical to those in Fig. 6f. See Fig. 6f for sample size and statistics. (d) Same as panel c, but comparing Single-shape trials to Character (stroke 2+) trials. Each data point is a unique pair of primitive-task conditions (trial-averaged). For testing primitive encoding, these pairs were the same task type, but different primitive [n = 858 (subject 1), 624 (subject 2)]. For task-type encoding, the pairs were the same primitive, but different task type [n = 90 (S1), 63 (S2). (e) Same as panel c, but comparing Single-shape trials to Multi-shape (stroke 2+) trials. For testing primitive encoding, these pairs were the same task type, but different primitive [n = 1458 (subject 1), 778 (subject 2)]. For task-type encoding, the pairs were the same primitive, but different task type [n = 117 (S1), 75 (S2)].

**Extended Data Fig. 15.**
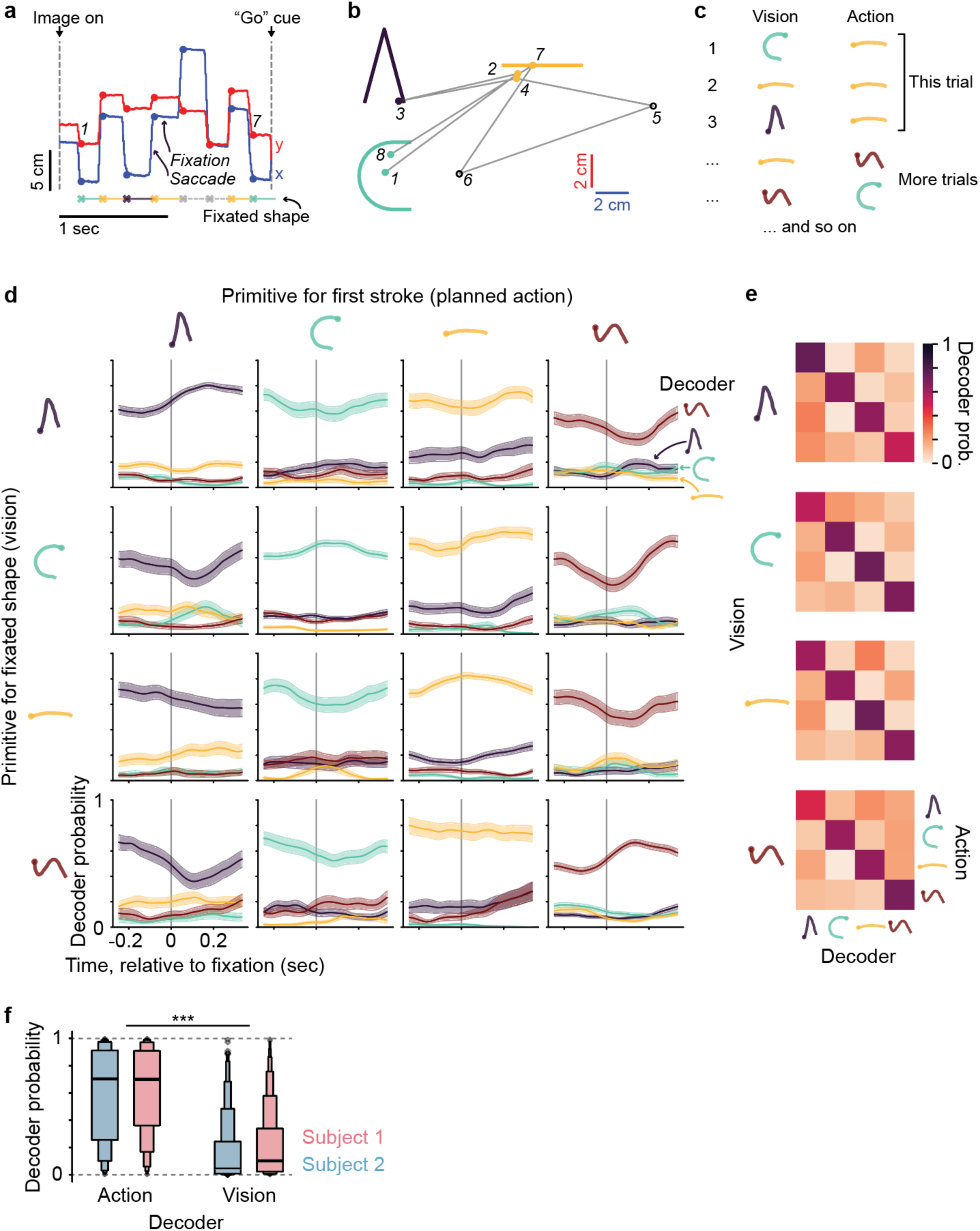
Primitive encoding in PMv reflects the first primitive the subject is planning to draw, rather than the primitive associated with the visually-fixated shape, in the Multi-shape task. (a) Gaze location during the planning epoch (between “image on” and “go cue”) for an example trial. Gaze involved interleaved saccades (rapid change in location) and fixations (stable location). The shape associated with each fixation is indicated by the colored line below (dashed gray means the fixation was too far from any shape and therefore not assigned any shape). On this trial, the image had three shapes (see panel b), and the subject chose to draw the horizontal line first. There were no imposed rules for what order to draw the shapes. (b) Saccades (grey lines) and fixations (circles) overlaid on the image for this example trial, numbered in order, and colored by the identity of the fixated shape. Note that shapes on the screen are colored here, but were not for the monkey. Each fixation event was assigned a “vision” label based on its closest image shape, and a “planned action” label based on which shape the subject chose to draw first on that trial. Fixations that were too far from any shape (here, numbers 5 and 6) were excluded from subsequent analyses (see Methods). (c) Table illustrating a dataset of fixation events collected across trials. Note the dissociation between “vision” and “action” labels across events. (d) Average fixation-aligned decoder probability for each primitive class, grouping fixation events by the planned action (columns) and the fixated shape (rows). This, and the following panels, includes fixation events from the second half of the planning epoch, when the subject is expected to have decided what to draw first. Note the high probability for the decoder for the planned action, and low probability for the decoder for the fixated shape. Further, note that the decoder probability for the planned action is not strongly modulated relative to the time of fixation onset. (e) Summary of mean decoder probability across conditions varying in vision and action. The mean is taken over the time window 0.05 to 0.3 sec after fixation onset. Note the strong diagonal band, consistent with activity encoding the planned primitive rather than the visually-fixated shape. (f) Summary of decoder probability for the action vs. vision, shown in a letter-value plot (see explanation in Extended Data Fig. 12b). Each fixation event contributed one data point for the planned action and one for the visually-fixated shape. This includes only fixation events with action differing from vision. ***, p < 0.0005 (each subject), Mann–Whitney U test, n = 883 fixations (subject 1) and n = 446 (subject 2), each from a single session.

**Extended Data Fig. 16.**
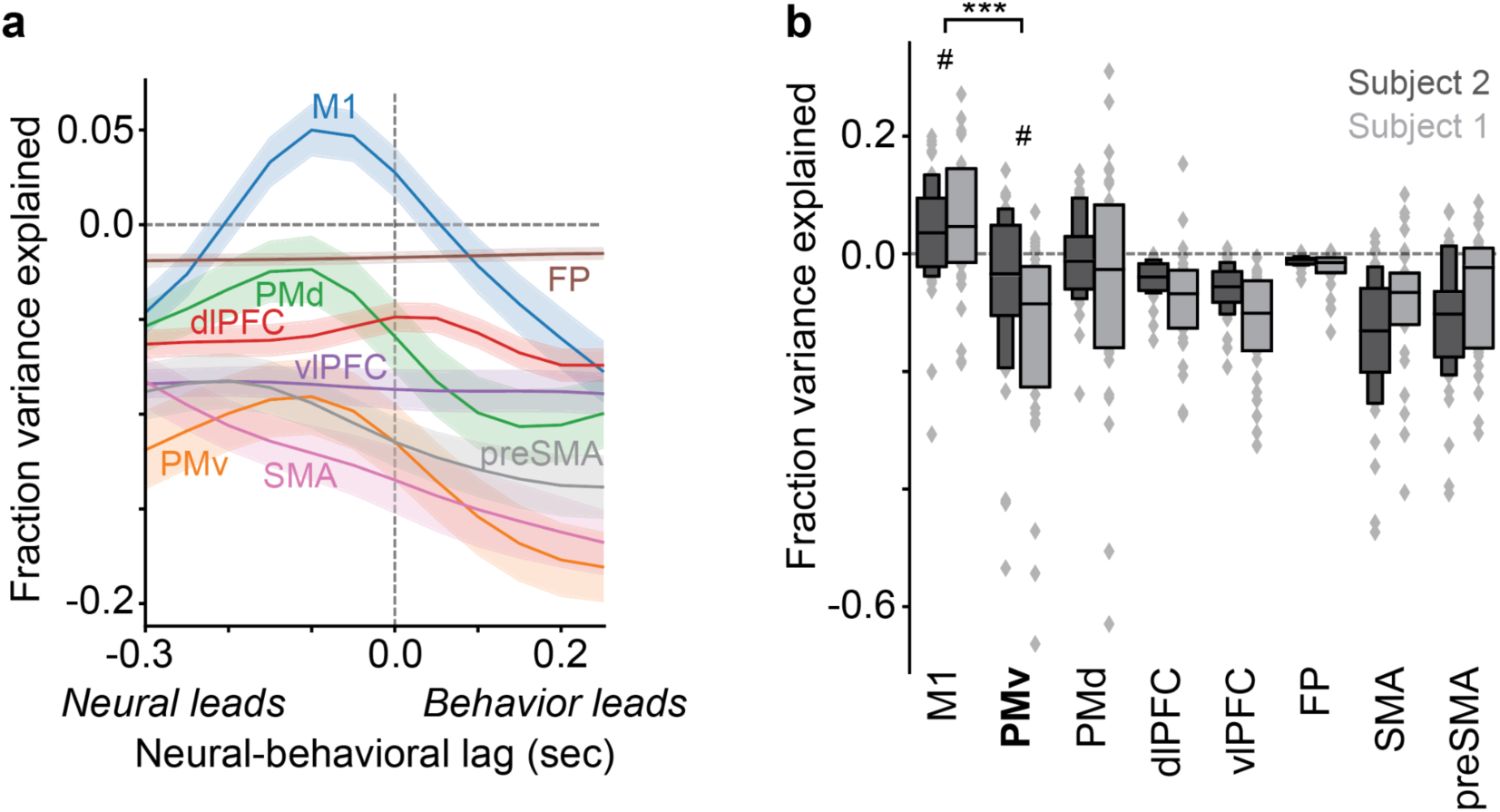
PMv activity does not significantly encode generalized motor kinematics. (a) Fraction of neural activity variance (across time bins and trials) explained by stroke kinematics (x- and y-velocity) during drawing. A linear regression model, mapping from kinematics to neural activity, was trained on a subset of data consisting of every primitive except one, and then tested on the one held-out primitive, therefore testing for encoding of kinematics that generalizes across primitives. This was repeated for every primitive (S1, n = 30 primitives across five sessions; S2, n = 42 primitives across three sessions; datasets were combined in this plot). This analysis was performed separately for different time lags between the neural and behavioral data, and the resulting values were plotted as a function of time lag. Note that negative explained variance indicates failure to generalize across primitives, with the model performing worse than an intercept-only model trained on the held-out dataset (see Methods). (b) Summary of fraction of variance explained, averaging over the time window -0.15 to -0.05 s relative to zero lag, summarized in a letter-value plot (see explanation in Extended Data Fig. 12b). Note that M1 is the only area with positive fraction variance explained. The M1 values (∼0.0 to 0.2) are comparable to a prior study of motor cortex in human handwriting^55^ (see Methods). Statistical tests involving M1 and PMv: ***, M1 vs. PMv: S1 (W = 10, p = 8.01 x 10^-8^), S2 (W = 137, p = 3.36 x 10^-5^), S1 and S2 combined (W = 205, p = 4.87 x 10^-10^). #, M1 vs. zero: S1 (W = 117, p = 0.016), S2 (W = 230, p = 0.0049), combined (W = 667, p = 0.00028), S1 does not reach significance after Bonferroni correction. #, PMv vs. zero: S1 (W = 33, p = 5.97 x 10^-6^), S2 (W = 293, p = 0.047), combined (W = 559, p = 2.27 x 10^-5^), S2 do not reach significance after Bonferroni correction. Sample size for all tests: S1, n = 30 primitives across five sessions; S2, n = 42 primitives across three sessions. Bonferroni correction considered all brain regions, thus correcting for 28 tests for PMv vs. M1, and for 8 tests for PMv/M1 vs. zero.

**Supplementary Video 1. Behavior in Character task (example character 1, subject 1).** This depicts the trial in Fig. 2l, 4th column from the right.

**Supplementary Video 2. Behavior in Character task (example character 1, subject 2).** This depicts the trial in Fig. 2l, 4th column from the right.

**Supplementary Video 3. Behavior in Character task (example character 2, subject 1).** This depicts the trial in Fig. 2l, 7th column from the left.

**Supplementary Video 4. Behavior in Character task (example character 2, subject 2).** This depicts the trial in Fig. 2l, 7th column from the left.

**Supplementary Video 5. Behavior in Character task (example character 3, subject 1).** This depicts the trial in Fig. 2l, 3rd column from the left.

**Supplementary Video 6. Behavior in Character task (example character 3, subject 2).** This depicts the trial in Fig. 2l, 3rd column from the left.

**Supplementary Video 7. Behavior in Character task (example character 4, subject 1).** This depicts the trial in Fig. 2l, 2nd column from the left.

**Supplementary Video 8. Behavior in Character task (example character 4, subject 2).** This depicts the trial in Fig. 2l, 2nd column from the left.

**Supplementary Video 9. Behavior in Character task (example character 5, subject 1).** This depicts the trial in Fig. 2l, 1st column from the left.

**Supplementary Video 10. Behavior in Character task (example character 5, subject 2).** This depicts the trial in Fig. 2l, 1st column from the left.

